# Co-transcriptional splicing regulates 3’ end cleavage during mammalian erythropoiesis

**DOI:** 10.1101/2020.02.11.944595

**Authors:** Kirsten A. Reimer, Claudia Mimoso, Karen Adelman, Karla M. Neugebauer

## Abstract

Pre-mRNA processing steps are tightly coordinated with transcription in many organisms. To determine how co-transcriptional splicing is integrated with transcription elongation and 3’ end formation in mammalian cells, we performed long-read sequencing of individual nascent RNAs and PRO-seq during mouse erythropoiesis. Splicing was not accompanied by transcriptional pausing and was detected when RNA polymerase II (Pol II) was within 75 – 300 nucleotides of 3’ splice sites (3’SSs), often during transcription of the downstream exon. Interestingly, several hundred introns displayed abundant splicing intermediates, suggesting that splicing delays can take place between the two catalytic steps. Overall, splicing efficiencies were correlated among introns within the same transcript, and intron retention was associated with inefficient 3’ end cleavage. Remarkably, a thalassemia patient-derived mutation introducing a cryptic 3’SS improves both splicing and 3’ end cleavage of individual β-globin transcripts, demonstrating functional coupling between the two co-transcriptional processes as a determinant of productive gene output.

## INTRODUCTION

Transcription and pre-mRNA processing steps - 5’ end capping, splicing, base modification, and 3’ end cleavage – required for eukaryotic gene expression are each carried out by macromolecular machines. The spliceosome assembles *de novo* on each intron, recognizing the 5’ and 3’ splice sites (SSs) that demarcate intron boundaries and then catalyzing two transesterification reactions to excise introns and ligate exons together (Wilkinson et al., 2020). In mammalian cells, genes typically encode pre-mRNAs containing 8-10 introns of variable lengths, creating a high cellular demand for spliceosomes relative to all of the other machineries, which only act once per transcript. Splicing is also a highly-regulated process; it is influenced by environmental factors, developmental cues, and factors in the local pre-messenger RNA (pre-mRNA) environment, such as RNA secondary structure and RNA-binding protein occupancy (Baralle and Giudice, 2017; Jeong, 2017; Lin et al., 2016; Pai and Luca, 2019). The influence of transacting factors on the selection of 5’ and 3’SSs is thought to explain how constitutive and alternative splice sites are chosen. These working models still largely rely on *in vitro* biochemistry and often do not explain changes in alternative splicing or overall gene expression observed upon experimental perturbation or disease-associated mutations of splicing factors (Joshi et al., 2017; Manning and Cooper, 2017). Thus, despite detailed knowledge of modulatory factors, the mechanisms underlying the gene regulatory potential of pre-mRNA splicing are not fully understood *in vivo.*

Across species, tissues, and cell types, splicing occurs during pre-mRNA synthesis by Pol II (Custodio and Carmo-Fonseca, 2016; Neugebauer, 2019). Thus, spliceosome assembly occurs as the nascent RNA is growing longer and more diverse in sequence and structure. Spliceosomes may not assemble on all of the introns at the same time, because promoter-proximal introns are synthesized before promoter-distal introns. The questions of whether introns are spliced in the order they are transcribed and how splicing of individual introns within a given transcript might be coordinated are currently the subject of intense investigation. Co-transcriptional splicing also demands that the constellation of splicing factors capable of regulating a splicing event bind the nascent RNA coordinately with the timing imposed by transcription and in a relevant spatial window. For example, a splicing inhibitor element in a given nascent RNA would only be influential if it were transcribed *before* the target intron was removed.

Recently, the Neugebauer lab has used single-molecule sequencing approaches to determine how splicing progresses as a function of transcription in budding and fission yeasts, where introns are removed shortly after synthesis (Alpert et al., 2020; Carrillo Oesterreich et al., 2016; Herzel et al., 2018). The approaches mark the nascent RNA’s 3’ end, which is present in the catalytic center of Pol II, to determine the position of Pol II when splicing occurs and define the sequence of the pre-mRNA substrate acted on by the spliceosome. These data show that only a small portion of the downstream exon may be needed for 3’SS identification and splicing. Interestingly, altering the rate of Pol II elongation affects splicing outcomes, including widespread changes in alternative splicing (Aslanzadeh et al., 2018; Braberg et al., 2013; Carrillo Oesterreich et al., 2016; de la Mata et al., 2003; Fong et al., 2014; Ip et al., 2011; Jonkers and Lis, 2015; Schor et al., 2013). Taken together, these findings suggest that transcription elongation rate may govern the amount of downstream RNA available for *cis* regulation at the time that splicing takes place. This in turn would determine which *trans*-acting regulatory factors could be recruited to the nascent RNA to modulate splicing. To obtain mechanistic insights into these processes, we need to understand how mammalian cells – with many more introns per gene and vastly increased levels of alternative splicing compared to yeast – coordinate co-transcriptional splicing with transcription elongation.

Another issue raised by co-transcriptional RNA processing is how splicing is coordinated with other pre-mRNA processing steps (Bentley, 2014; Herzel et al., 2017). In recent long-read sequencing studies in budding and fission yeasts (Alpert et al., 2020; Herzel et al., 2018), “all or none” splicing of individual nascent transcripts was discovered, suggesting positive and negative cooperativity among neighboring introns and polyA cleavage sites. Indeed, crosstalk among introns was observed in human cells at the same time by others (Kim et al., 2017; Tilgner et al., 2018). However, those studies did not explore coupling to 3’ end formation. Cleavage of the nascent RNA by the cleavage and polyadenylation machinery at polyA sites (PAS) releases the RNA from Pol II and the RNA is subsequently polyadenylated (Kumar et al., 2019). Coupling between splicing and 3’ end cleavage is important, because uncleaved transcripts are degraded by the nuclear exosome in *S. pombe* (Herzel et al., 2018; Meola et al., 2016; Zhou et al., 2015). Whether 3’ end cleavage efficiency contributes to gene expression levels in mammalian cells is currently unknown.

Here we report our analysis of nascent RNA transcription and splicing in murine erythroleukemia (MEL) cells undergoing erythroid differentiation, a developmental program that exhibits well-known, drastic changes in gene expression (An et al., 2014; Reimer and Neugebauer, 2018). We have employed two single-molecule sequencing approaches to directly measure co-transcriptional splicing of nascent RNA: *(i)* Long-read sequencing (LRS), which enables genome-wide analysis of splicing with respect to Pol II position and *(ii)* Precision Run-On sequencing (PRO-seq), enabling the assessment of Pol II density at these sites. We rigorously determine the spatial window in which co-transcriptional splicing occurs and define co-transcriptional splicing efficiency for thousands of mouse introns, Pol II elongation behavior across splice junctions, and the effects of efficient co-transcriptional splicing on 3’ end cleavage. These findings identify the pre-mRNA substrates of splicing and show that splicing of multiple introns within individual transcripts is coordinated with 3’ end cleavage. In particular, the demonstration of highly efficient splicing in the absence of transcriptional pausing causes us to rethink key features of splicing regulation in mammalian cells.

### RESULTS

#### PacBio Long-read Sequencing of Nascent RNA Yields High Read Coverage

Murine erythroleukemia (MEL) cells are immortalized at the proerythroblast stage and can be induced to enter terminal erythroid differentiation by treatment with 2% DMSO for five days (Antoniou, 1991). Phenotypic changes include decreased cell volume, increased levels of β-globin, and visible hemoglobinization (**Figures S1A-C**). We used chromatin purification of uninduced and induced MEL cells to enrich for nascent RNA (**Figure 1A**). Chromatin purification under stringent washing conditions allows release of contaminating RNAs and retains the stable ternary complex formed by elongating Pol II, DNA, and nascent RNA (**Figure S1D**; (Wuarin and Schibler, 1994). Importantly, spliceosome assembly does not continue during chromatin fractionation or RNA isolation, because the presence of the splicing inhibitor Pladienolide B throughout the purification process does not change splicing levels (**Figure S2**).

**Figure 1.**
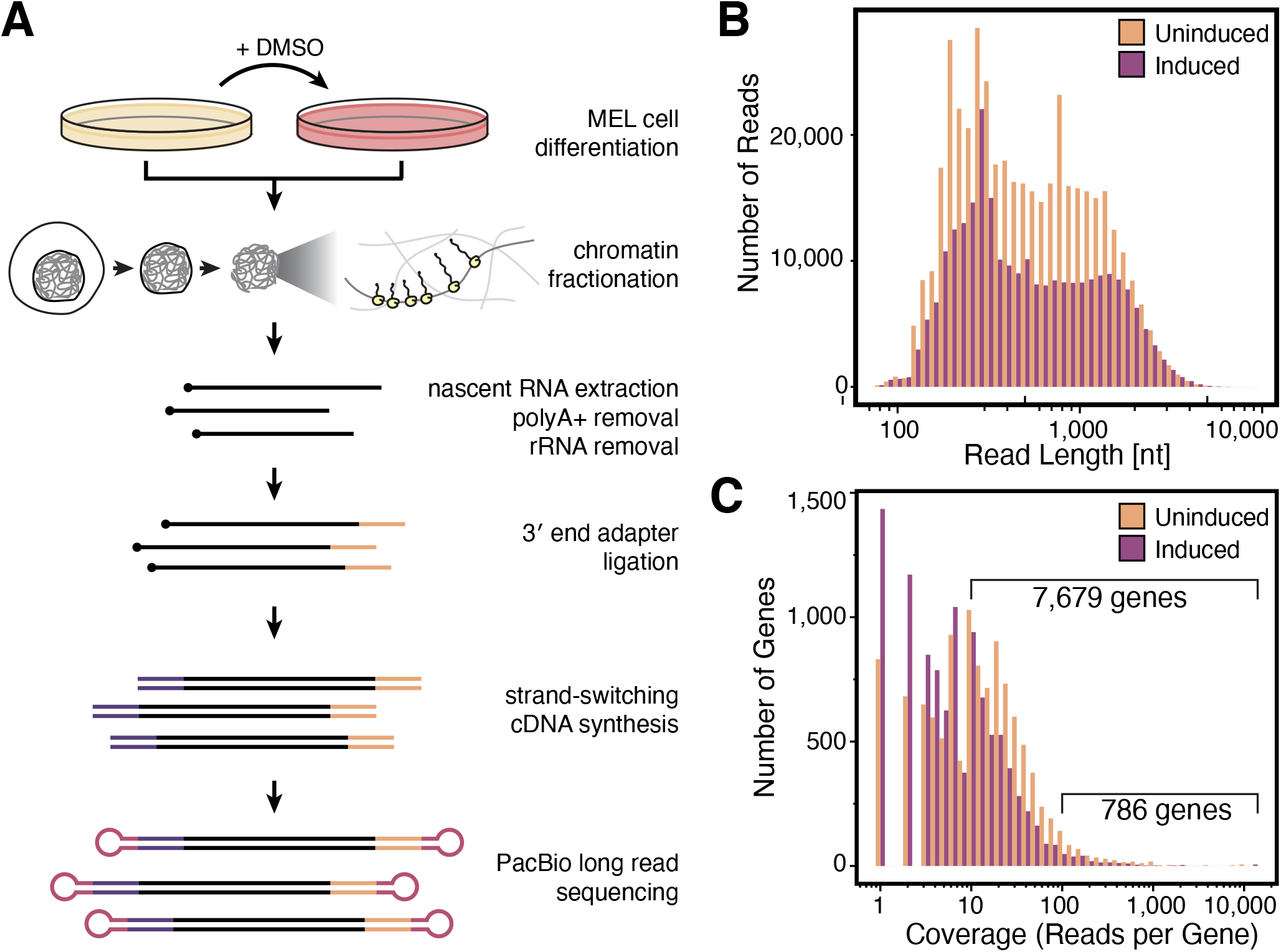
Long-read sequencing of nascent RNA from differentiating mouse erythroblasts. **(A)** Schematic of nascent RNA isolation and sequencing library generation. MEL cells are treated with 2% DMSO to induce erythroid differentiation, cells are fractionated to purify chromatin, and chromatin-associated nascent RNA is depleted of polyadenylated and ribosomal RNAs. An adapter is ligated to the 3’ ends of remaining RNAs, then a strand-switching reverse transcriptase is used to create doublestranded cDNA that is the input for PacBio library preparation. **(B)** Read length and **(C)** read depth distribution of PacBio long-reads. See also **Figures S1** and **S2**, and **Table S1**.

To generate libraries for LRS, we established the protocol outlined in **Figure 1A**. Two biological replicates, each with two technical replicates, were sequenced using PacBio RSII and Sequel flow cells, yielding a total of 1,155,629 mappable reads (**Table S1**). Reads containing a non-templated polyA tail comprised only 1.7% of the total reads (**Table S1**) and were removed bioinformatically along with abundant 7SK RNA reads. Of the remaining reads, the average read length was 710 and 733 nucleotides (nt), and the average coverage in reads per gene was 8.4 and 4.8 for uninduced and induced samples, respectively (**Figure 1B-C**). More than 7,500 genes were represented by more than 10 reads per gene in each condition (**Figure 1C**). Coverage of 5’ ends was focused at annotated transcription start sites (TSSs), with 18.3% of 5’ ends within 50 bp of an active TSS across all samples. As expected, 3’ end coverage was distributed more evenly throughout gene bodies, with an increase just upstream of annotated transcription end sites (TESs) and a drop after TESs (**Figure S1E**).

#### LRS Reveals Rapid and Efficient Co-transcriptional Splicing

Each long-read provides two critical pieces of information: the 3’ end reveals the position of Pol II when the RNA was isolated; the splice junctions reveal if splicing has occurred and which splice sites were chosen. Here, we present our LRS data in a format that highlights 3’ end position and the associated splicing status (**Figure 2A&B**; **Figure S3A**). Each transcript was categorized and colored according to its splicing status, which can be either “all spliced”, “partially spliced”, “all unspliced”, or “NA” (transcripts that did not span an entire intron or a 3’SS). For each gene, we calculated the fraction of long-reads that were all spliced, partially spliced, or all unspliced (**Figure 2A**; bar plot far right), enabling a survey of splicing behaviors within individual transcripts (Alpert et al., 2020; Herzel et al., 2018; Kim et al., 2017).

**Figure 2.**
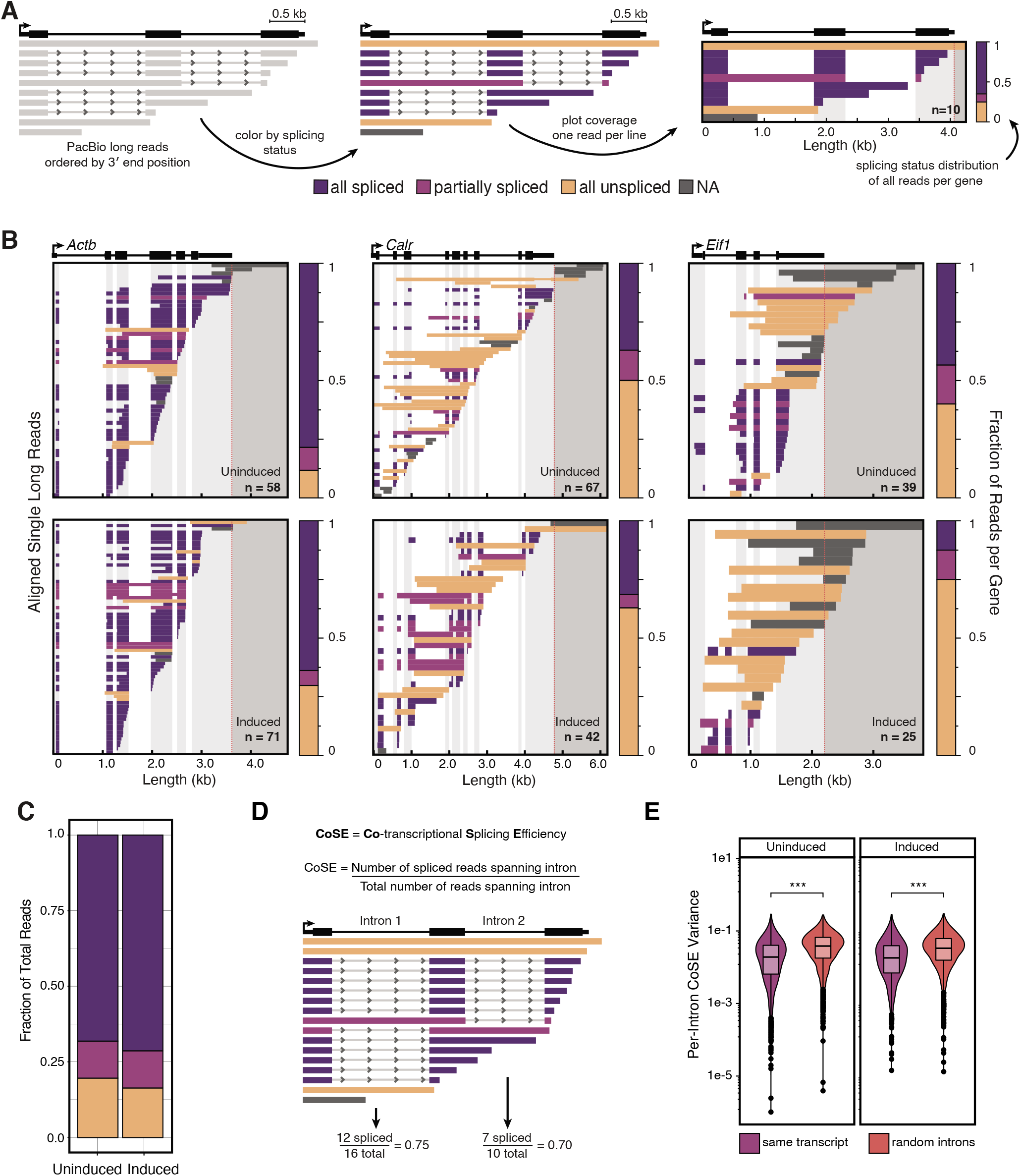
Individual mammalian nascent RNA sequences reveal coordination of co-transcriptional splicing. **(A)** LRS data visualization for analysis of co-transcriptional splicing. Gene diagram is shown at the top, with the black arrow indicating the TSS. Reads are aligned to the genome and ordered by 3’ end position. Color code indicates the splicing status of each transcript. Each horizontal row represents one read. Panels at far right and below: regions of missing sequence (e.g. spliced introns) are transparent. Light gray shading indicates regions of exons, and dark gray shading indicates the region downstream of the annotated PAS (dotted red line). The number of individual long-reads aligned to each gene (n) is indicated. Bar graph at the far right of each plot indicates the fraction of reads that are all spliced (dark purple), partially spliced (light purple), or all unspliced (yellow) for that gene. **(B)** LRS data are shown for uninduced (top) and induced (bottom) MEL cells for three representative genes: *Actb, Calr, Eif1.* **(C)** Fraction of long-reads that are all spliced, partially spliced, or all unspliced (n = 120,143 reads uninduced, n = 71,639 reads induced). **(D)**. For each intron that is covered by 10 or more reads, CoSE is defined as the number of reads that are spliced divided by the total number of reads that span the intron. **(E)** Variance in CoSE for transcripts that include 3 or more introns covered by at least 10 reads (n = 1,240 transcripts uninduced, n = 788 transcripts induced) compared to the variance in CoSE for a randomly selected group of introns. Significance tested by Mann Whitney U-test; *** represents p-value < 0.001. See also Figure S3.

Splicing status of individual transcripts varied from gene to gene. For example, the gene *Actb* had mostly all spliced reads (78% and 75% of reads in uninduced and induced cells respectively), while *Calr* and *Eif1* had a greater fraction of all unspliced reads (**Figure 2B**). Genome-wide, the majority of long-reads were all spliced (**Figure 2C**; 68.0% and 73.8% for uninduced and induced cells, respectively), with an average of 88% of all introns being spliced. Therefore, the majority of introns are removed co-transcriptionally. To validate this finding, we examined the read length distribution for reads of each splicing status (**Figure S3B**). As expected, partially spliced and all unspliced reads were longer than all spliced reads due to the presence of introns, suggesting that the efficient shortening of nascent RNA due to splicing limits the lengths of long-reads.

To quantify co-transcriptional splicing for each intron detected by at least 10 long-reads, we defined a metric termed the <u>Co</u>-transcriptional <u>S</u>plicing <u>E</u>fficiency (CoSE), tabulated as the number of spliced reads that span the intron divided by the total number of reads (spliced + unspliced) that span the intron (**Figure 2D)**. A higher CoSE value indicates a higher fraction of co-transcriptional splicing. To validate this metric, we analyzed an independently generated total RNA-seq dataset in uninduced MEL cells (downloaded from ENCODE; (Davis et al., 2018)). Although nascent RNA is rare in total RNA, the density of reads mapping to a given intron is expected to be inversely proportional to splicing efficiency. The ratio of intron-mapping reads relative to the flanking exon-mapping reads was calculated for each intron and compared to CoSE levels. As expected, higher CoSE corresponded to lower relative intron coverage in the total-RNA seq data (**Figure S3C**). Thus, this independent data set validates the CoSE metric. CoSE values also remained stable across all levels of read coverage (**Figure S3D**).

To determine if intron splicing events are coordinated within the same transcript, we asked how similar CoSE values were between introns in the same transcript. To do so, transcripts containing at least 3 introns with recorded CoSE values (n = 2,028) were compiled. We found that the variance in CoSE between introns within the same transcript was significantly smaller than the variance in CoSE for the same number of randomly assorted introns (**Figure 2E**); these differences persisted when we analyzed transcripts containing 3, 4, or 5 introns supported by long-reads (**Figure S3E**). Taken together, these results suggest that most introns are well-spliced co-transcriptionally, and that splicing is coordinated in mammalian multi-intron transcripts expressed by both uninduced and induced MEL cells.

The frequency of all-spliced nascent transcripts implies that splicing in mammalian cells is rapid enough to match the rate of transcription. A direct way to address this is to measure the position of Pol II on nascent RNA when ligated exons are observed. Observing Pol II downstream of a spliced junction indicates that the active spliceosome has assembled and catalyzed splicing in the time it took for Pol II to translocate the measured distance. Therefore, we determined the distance in nucleotides between the 3’ end of each read and the nearest spliced exon-exon junction (**Figure 3A**). To eliminate 3’ ends that arise from splicing intermediates and not from active transcription, reads with 3’ ends mapping precisely to the last nt of exons were removed from this analysis. Although the longest distances between splice junctions and elongating Pol II were just over 6 kb, these were rare. Instead, 75% of splice junctions were within ~300 nt of a 3’ end, and the median distance was 154 nt in uninduced cells and 128 nt in induced cells (**Figure 3B**) Therefore, changes in the gene expression program during erythropoiesis did not alter the dynamic relationship between transcription and splicing. Consistent with this, CoSE values were similar when comparing induced to uninduced cells (**Figure 3C**; Spearman’s rho = 0.56). In fact, only 66 introns with improved splicing, and 42 introns with reduced splicing displayed > 2-fold change in CoSE upon induction. Taken together, this analysis shows that although global changes in gene expression take place between these two timepoints, the relationship between transcription and splicing remains the same. Overall, these two measurements do not support major changes in splicing efficiency during erythroid differentiation. Moreover, the distance from Pol II to the nearest splice junction was independent of GO category or intron length (**Figure S4B**; GO analysis not shown). Because median exon size in the mouse genome is 151 nt (Waterston et al., 2002), our data indicate that active spliceosomes can be fully assembled and functional when Pol II is within or just downstream of the next transcribed exon. Recent direct sequencing of nascent RNA seemed to reveal less rapid splicing (Drexler et al., 2020). However, when we analyzed this dataset in the same manner as our own, the cumulative distance from Pol II to the nearest splice junction is similarly close across organisms and cell types (median distance in human BL1184 = 244 nt, human K562 = 310 nt, *Drosophila* S2 = 335 nt; **Figure S4A**). Thus, we conclude that efficient and coordinated splicing are a general property of metazoan gene expression.

**Figure 3.**
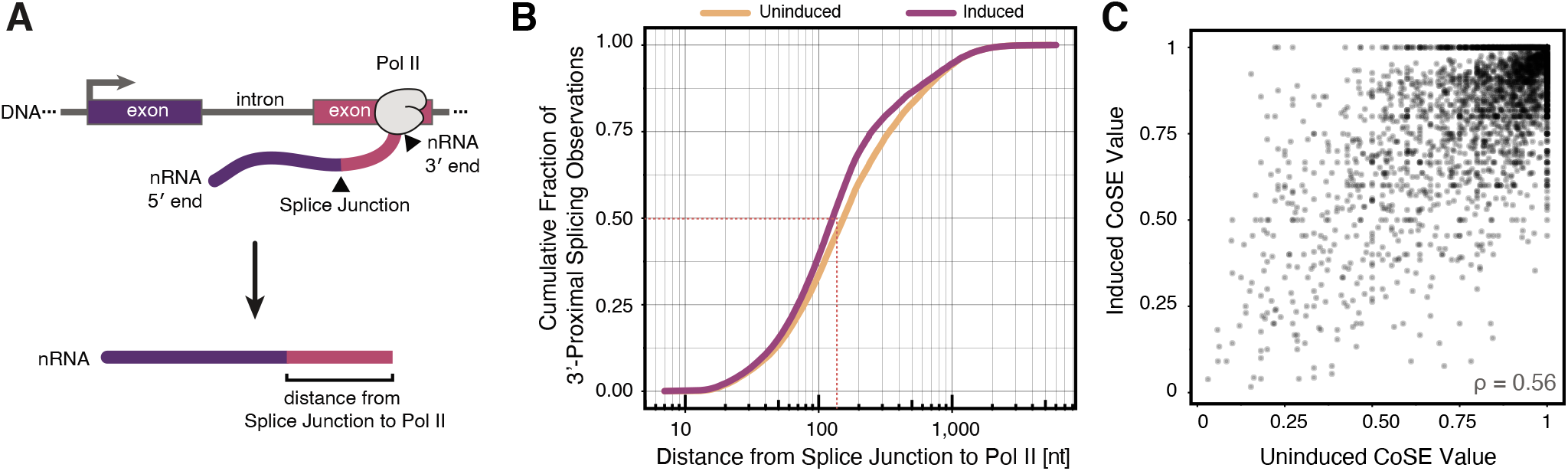
Spliceosome-Pol II proximity is unchanged by differentiation. **(A)** Schematic definition of the distance from the 3’ end of a nascent RNA (nRNA) to the most 3’-proximal splice junction. 3’ end sequence reports the position of Pol II when nascent RNA was isolated. **(B)** Distance (nt) from the 3’-most splice junction to Pol II position is shown as a cumulative fraction for uninduced and induced cells (n = 101,911 observations uninduced, n = 66,656 induced). **(C)** CoSE in induced and uninduced conditions. Each point represents a single intron which is covered by at least 10 long-reads in both induced and uninduced conditions. Spearman’s rho = 0.56, n = 4,170 introns. See also Figure S4.

#### Pol II Does Not Pause at Splice Sites for Splicing to Complete

One explanation for the relatively short distances observed between splice junctions and Pol II may be that Pol II pauses just downstream of an intron, allowing time for splicing to occur before elongation continues. Alternatively, Pol II could pause at the end of the downstream exon. This model has been proposed as a mechanism for splicing and transcription to feedback on each other, with pausing providing a possible checkpoint for correct RNA processing (Alexander et al., 2010b; Carrillo Oesterreich et al., 2011; Chathoth et al., 2014; Milligan et al., 2017). However, recent work has disagreed on the behavior of Pol II elongation near splice junctions, with some studies indicating long-lived pausing at splice sites, and others reporting no significant pausing (Kwak et al., 2013; Mayer et al., 2015; Sheridan et al., 2019). To resolve this controversy, we measured changes in elongating Pol II density genome-wide using Precision Run-On sequencing (PRO-seq) in MEL cells. PRO-seq maps actively elongating Pol II complexes at single-nucleotide resolution by incorporating a single biotinylated NTP (Mahat et al., 2016). Comparing PRO-seq with LRS is advantageous, because PRO-seq data provide an independent measure of nascent RNA 3’ ends that are undergoing active elongation; these 3’ ends cannot originate from other chromatin-associated intermediates, such as splicing intermediates (see below).

We analyzed our PRO-seq data to determine if transcription elongation behavior changes across intron-exon boundaries. Because both induced and uninduced LRS datasets showed an overlapping distribution of Pol II when spliced products are observed, we initially combined the PRO-seq datasets. As expected, metagene plots around active TSSs revealed prominent promoter-proximal pausing (**Figure 4A**; (Core and Adelman, 2019). Analyzing PRO-seq signal around splice sites initially revealed a small peak near the 5’SS. To control for the possibility that high PRO-seq density from TSS peaks might bleed through to the first 5’SS, first introns were independently analyzed. Indeed, elevated PRO-seq signal in the vicinity of 5’SSs was only seen at first introns, and only at introns with 5’SS ≤ 250 nt from the TSS (**Figure S5A**). Interestingly, middle introns showed a similar profile to terminal introns, both lacking increased PRO-seq signal around splice sites (**Figure S5B**). Accordingly, after removal of first introns from our analysis, PRO-seq signal showed only minor fluctuations around 5’SSs and 3’SSs (**Figure 4A**).

**Figure 4.**
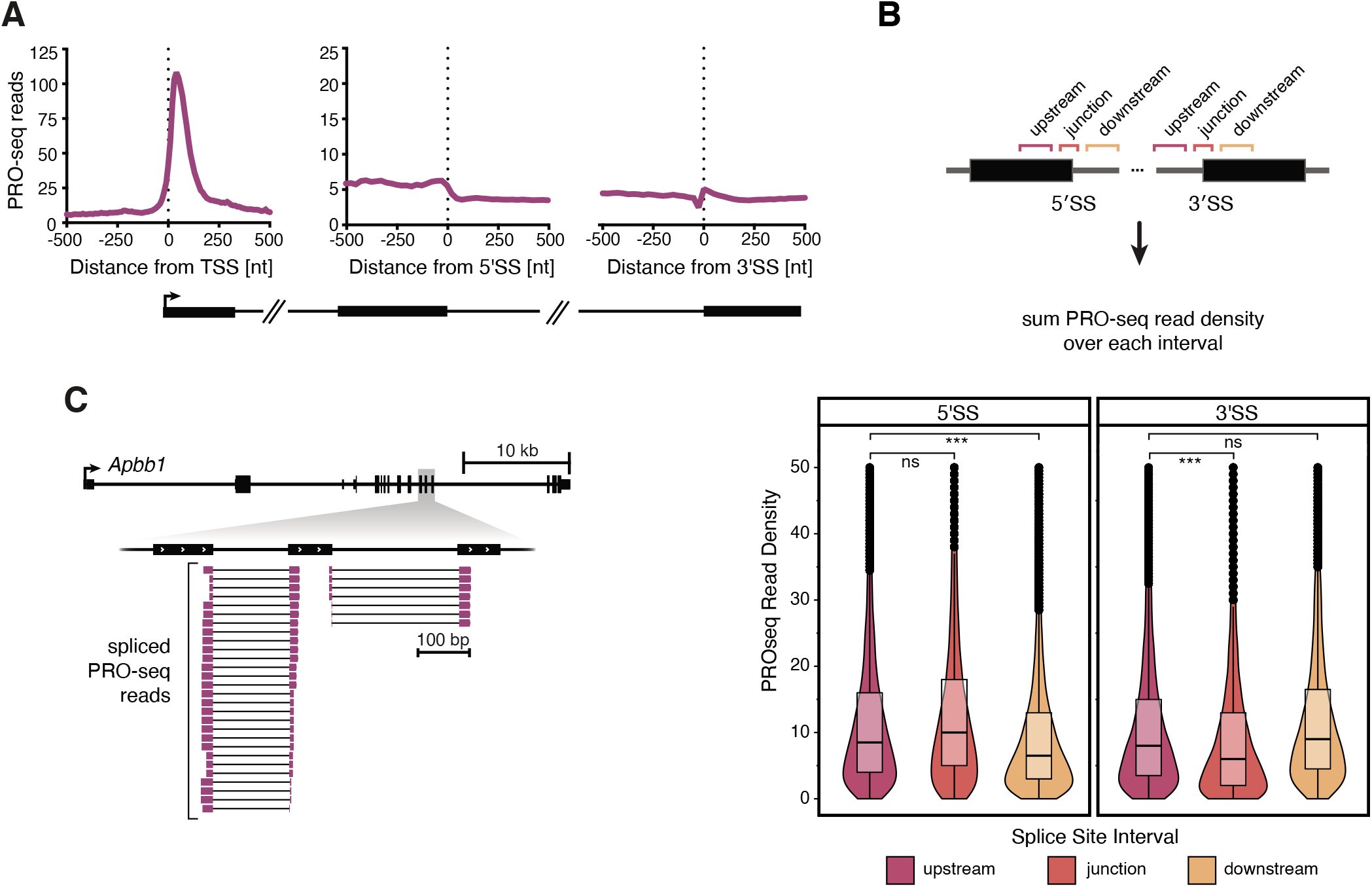
Pol II does not pause at 5’ or 3’ splice sites. **(A)** PRO-seq 3’ end coverage is shown aligned to active transcription start sites (TSS), 5’ splice sites (5’SS), and 3’ splice sites (3’SS). **(B)** Top: Schematic illustrating the use of color-coded intervals to quantify PRO-seq reads around each 5’SS and 3’SS to test for significance of pausing. Bottom: PRO-seq read density summed in each of the intervals indicated above around 5’SSs (left) and 3’SSs (right) from introns with at least 10 reads in uninduced conditions (n = 3,505). Significance tested by paired t-test; *** represents p-value < 0.001, ns represents p-value > 0.05. **(C)** Genome browser view showing spliced PRO-seq reads aligned to the *Apbb1* gene, where 3’ ends of reads represent the position of elongating Pol II. Only spliced reads, filtered from all reads, are shown. See also **Figure S5**.

The depth of our PRO-seq libraries enabled us to determine if these minor fluctuations in PRO-seq signal were statistically significant. To do so, we compared summed PRO-seq read counts in three windows surrounding each intron/exon junction for uninduced cells only (**Figure 4B**): −150 to −50 nt upstream of the junction, −40 to +10 nt spanning the junction, and +50 to +150 nt downstream of the junction. Comparing the signal between the upstream window and 5’SS-spanning window indicated no statistically significant increase in PRO-seq read density (paired t test p > 0.05). In contrast, a significant decrease in PRO-seq signal is observed as Pol II moves from the exon into the intron (p < 0.0001; comparing upstream to downstream window at the 5’SS), consistent with data showing faster transcription rates within introns (Jonkers et al., 2014; Veloso et al., 2014). Interestingly, we observe a dip in PRO-seq signal right before the 3’SS (p < 0.0001), instead of a peak of Pol II consistent with pausing. Although the cause of this dip remains to be determined, it is unlikely to represent Pol II arrest and/or termination near this junction because signal downstream of the 3’SS does not decrease. In summary, we can detect significant changes in Pol II elongation behavior as it moves from exon to intron and *vice versa,* but we find no evidence for significant pausing at splice junctions.

To rigorously compare splicing efficiency to Pol II elongation behavior, we evaluated PRO-seq signals around introns binned by CoSE values from our LRS data. Again, we observed no significant differences in Pol II profile around splice sites within any group of introns (**Figure S5C**). Interestingly, the overall level of PRO-seq coverage is lower in transcripts with higher CoSE values, suggesting that lowly expressed transcripts might be more efficiently spliced. Finally, a number of PRO-seq reads contained spliced junctions despite the short-read lengths (**Figure 4C**; 396,257 spliced reads out of 289,610,781 total mapped reads). These data confirm that mammalian splicing can occur when actively engaged Pol II is just downstream of the 3’SS, within a median distance of 128-154 nt. Taken together, two complementary methods to probe Pol II position and splicing status indicate that splicing can occur when Pol II is in close proximity to the 3’SS and in the absence of transcriptional pausing.

#### Splicing Intermediates are Abundant for a Subset of Introns

We expected to readily observe transient splicing intermediates in our nascent RNA libraries, because the two-step transesterification reaction yields a 3’-OH at the end of the upstream exon after step I (**Figure 5A**). Indeed, splicing intermediates have previously been observed using other chromatin-associated RNA sequencing methods (Burke et al., 2018; Chen et al., 2018; Churchman and Weissman, 2011; Nojima et al., 2015; Nojima et al., 2018). As expected, we observed elevated 3’ end coverage precisely at the last nucleotide of exons (**Figure 5B**), with these first step splicing intermediate reads accounting for 7.0% of the data (**Table S1**). The rarity of splicing intermediates detected agrees with our finding that splicing does not continue during chromatin fractionation or RNA isolation (**Figure S2**). Nevertheless, a small number of genes contained a large number of splicing intermediates at a single intron within the gene (**Figure 5C**; **Figure S6A**). For example, 216 of the 433 reads mapped to *Alas2* had 3’ ends mapped to the end of exon four (**Figure 5D**). We note that several reads (16/433) mapping to *Alas2* were one of the extremely rare instances of potential recursive splicing. In this case, we observed an unannotated splice junction which generated a new 5’SS immediately adjacent, with the junction sequence cagGUAUGU (**Figure 5E**). While we observed no other compelling instances of recursive splicing in our dataset, we note that recursive splicing has been previously characterized in extremely long introns, which are difficult to detect using our methods (Pai et al., 2018; Sibley et al., 2015). Nevertheless, we conclude that recursive splicing could occur co-transcriptionally.

**Figure 5.**
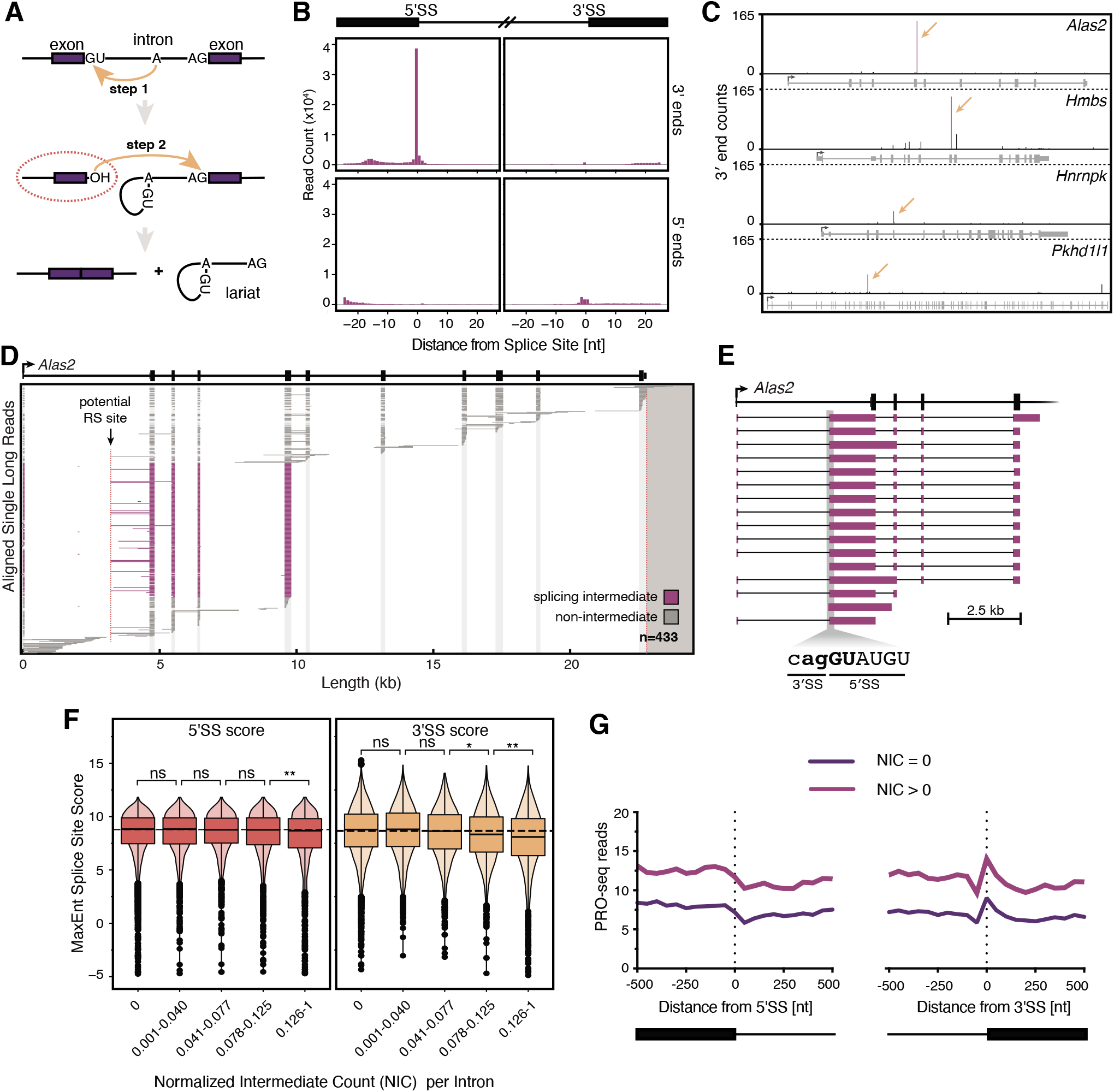
Splicing intermediates are abundant at introns with weak 3’ splice sites. **(A)** Schematic definition of first step splicing intermediates (dotted red oval), which have undergone the first step of splicing and have a free 3’-OH that can be ligated to the 3’ end DNA adapter. Splicing intermediate reads are characterized by a 3’ end at the last nucleotide of the upstream exon. **(B)** Coverage of long-read 3’ ends (top panels) and 5’ ends (bottom panels) aligned to 5’SSs (left) and 3’SSs (right) of introns. **(C)** Coverage of long-read 3’ ends across four example genes. Arrows indicate the positions where the most abundant splicing intermediates are observed. **(D)** Individual long-reads are shown for the gene *Alas2*. Diagram is similar to **Figure 2**, but individual reads are colored depending on whether they are splicing intermediates (purple) or not (gray). Data for uninduced and induced cells are shown combined. Potential recursive splicing site is indicated by an arrow and dotted line; recursively spliced reads are shown in detail in **(E)**. **(F)** MaxEnt splice site scores for 5’SS (left) and 3’SS (right) for introns with a coverage of at least 10 long-reads is shown categorized by the normalized intermediate count (NIC) at each intron. Introns with NIC = 0 (n = 3,890) are shown separately, and all other introns with NIC > 0 (n = 2,647) are separated in quartiles with NIC values shown. **(G)** Raw PRO-seq 3’ end coverage from uninduced cells aligned to 5’SSs, and 3’SSs for introns with NIC = 0 (n = 4,402), or NIC > 0 (n = 3,427). See also Figure S6.

To determine what features of specific introns might lead to increased splicing intermediates, we counted and normalized the number of splicing intermediates observed for each intron. The normalized intermediate count (NIC), is defined as the number of splicing intermediate reads at the last nt of the exon divided by the sum of splicing intermediate reads and spliced reads. This metric reports the fraction of long-reads that are captured between step I and step II of splicing. All unique introns which were covered by at least 10 long-reads were binned based on their observed NIC value, and the splice site strength of the introns in each bin was calculated using the MaxEnt algorithm (Yeo and Burge, 2004). While the 5’SS score was relatively constant, introns with the highest NIC value tended to have lower 5’SS and 3’SS scores (**Figure 5F**). Intron length or GC-content showed no similar trend (**Figure S6B-C**). Finally, we tested if spliceosomal stalling between steps I and II could be associated with Pol II pausing, by analyzing PRO-seq signals downstream of the associated 3’SSs. Note that PRO-seq signal was universally higher in introns with 1 or more splicing intermediates as was total RNA-seq density in flanking exons, indicating that these genes were more highly expressed (**Figure S6D-E**). However, no differences in Pol II density were detected around 3’SS between introns with NIC = 0 and NIC ? 1 (**Figure 5G**). Based on these data, we suggest that introns with weak 3’SSs experience a delay between the catalytic steps of splicing without a delay in transcription.

#### Unspliced Transcripts Display Poor Cleavage at Gene Ends

Consistent with physiological terminal erythroid differentiation, our induced MEL cells shifted to maximal expression of a- and β-globin genes, each containing two introns. Markedly increased numbers of long-reads mapped to the β-globin (*Hbb-b1*) locus were detected, in agreement with increased β-globin mRNA levels (**Figure 6A**; **Figure S1C**). To our surprise, a large fraction of individual β-globin long-reads in the induced condition had 3’ ends that were up to 2.5 kb downstream of the annotated polyA site (PAS), indicating that these transcripts failed to undergo 3’ end cleavage at the PAS. Pol II occupancy past the β-globin PAS was confirmed by PRO-seq (**Figure 6B**). Notably, PRO-seq reads are commonly detected well past the gene 3’ ends due to transcription termination (Core et al., 2008). However, our LRS data indicate significant transcription past the polyA cleavage site in the absence of 3’ end cleavage, which cannot be revealed by PRO-seq. Remarkably, the β-globin transcripts that escaped 3’ end cleavage were almost uniformly unspliced (**Figure 6A**). The α-globin genes (*Hba-a1* and *Hba-a2*) displayed a similar phenomenon (**Figure S7A-C**). Thus, a significant fraction of nascent globin RNAs undergo “all or none” RNA processing under these conditions of erythroid differentiation: either both introns are efficiently spliced and the nascent RNA is cleaved at the 3’ end or, conversely, both introns are retained and the nascent RNA is inefficiently cleaved.

**Figure 6.**
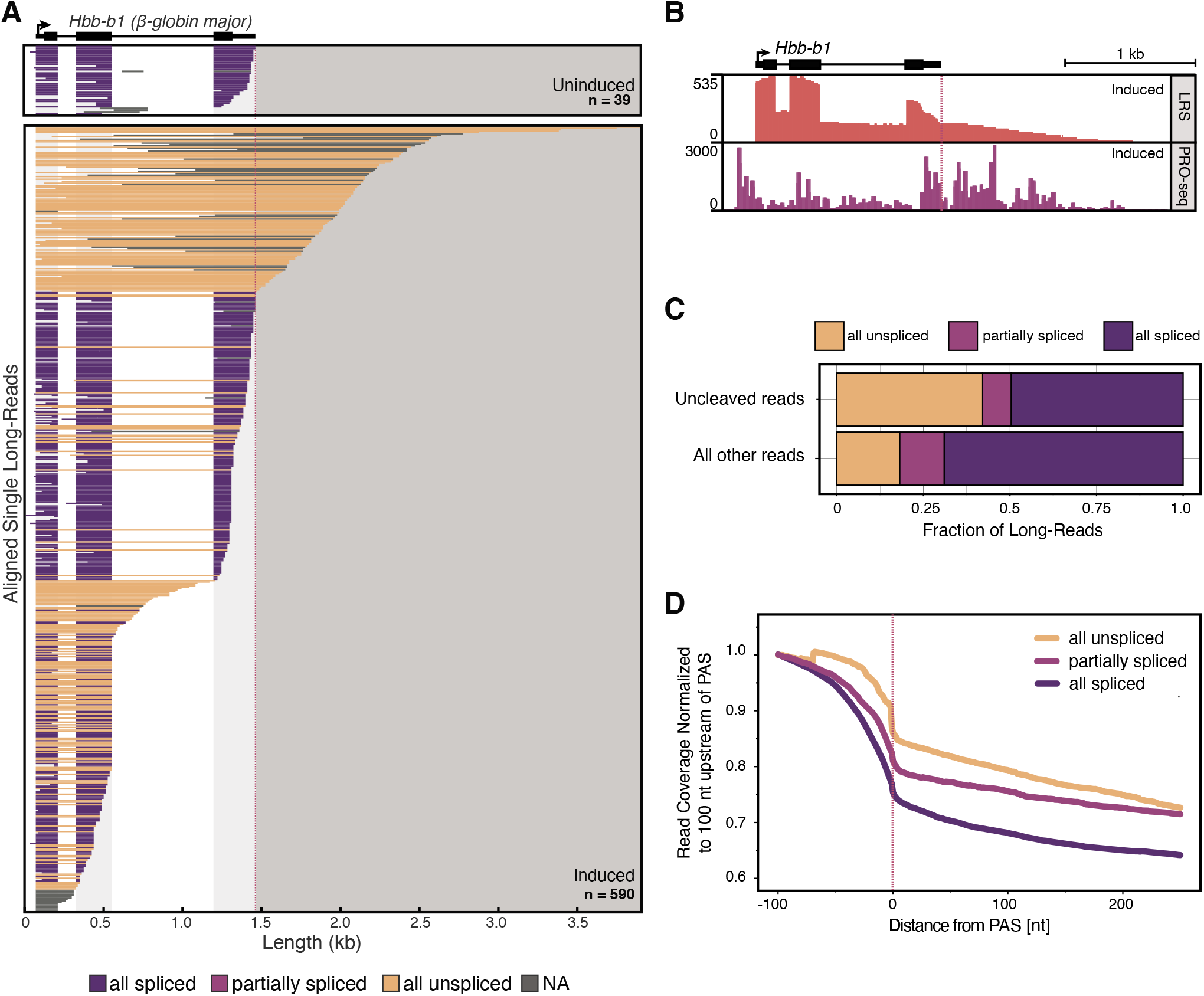
Poor splicing efficiency is associated with inefficient 3’ end cleavage. **(A)** Individual long-reads are shown for the major β-globin gene (*Hbb-b1*). Diagram is as described in **Figure 2**. **(B)** LRS coverage (orange) and PRO-seq 3’ end coverage (purple) in induced cells is shown at the *Hbb-b1* gene. Scale at the left indicates coverage in number of reads, and red dotted line indicates PAS. We note that the duplicated copies of β-globin in the genome (*Hbb-b1* and *Hbb-b2*) impedes unique mapping of short PRO-seq reads in the coding sequence, artificially reducing gene body reads. **(C)** Fraction of uncleaved long-reads (top) and all other long-reads (bottom) categorized by splicing status (as described in **Figure 2**). Uncleaved reads have a 5’ end within an actively transcribed gene region and a 3’ end greater than 50 nt downstream of the PAS (n = 5,694 uncleaved long-reads, and n = 172,612 other long-reads). **(D)** Long-read coverage in the region downstream of PASs is shown for long-reads separated by their splicing status. Coverage is normalized to the position 100 nt upstream of each PAS (n = 35,982 all unspliced reads, n = 24,102 partially spliced reads, and n = 134,581 all spliced reads). Red dotted line indicates PAS position. See also **Figure S7**.

Because other genes showed evidence of all-or-none splicing (**Figure 2B**; **Figure S3A**), a correlation between splicing and cleavage at the PAS was examined globally. To do so, we categorized long-reads as uncleaved if the 5’ end originated within a gene body and the 3’ end mapped more than 50 nt downstream of an annotated PAS. In comparison to all other long-reads, uncleaved long-reads were 2.5-fold more likely to be unspliced (**Figure 6C**). Next, we analyzed long-read coverage downstream of annotated PASs for reads with different splicing statuses. Coverage of all unspliced reads was globally higher in the region downstream of a PAS than it was for partially spliced or all spliced reads (**Figure 6D**). PRO-seq data support this claim, with significantly less PRO-seq signal observed in the region downstream of the PAS for transcripts harboring introns with the highest CoSE values (**Figure S7D**). This genome-wide decrease in splicing efficiency associated with impaired 3’ end cleavage confirmed the coordination between splicing and 3’ end processing prominently observed in the globin genes.

#### A β-thalassemia Mutation Enhances Splicing and 3’ End Cleavage Efficiencies

To investigate how mutations in splice sites alter co-transcriptional splicing efficiency, we took advantage of a known β-thalassemia allele. A patient-derived G>A mutation in intron 1 of human β-globin (*HBB*) leads to new AG dinucleotide in intron 1, creating a cryptic 3’SS 19 nt upstream of the canonical 3’SS (**Figure 7A**). This thalassemia-causing mutation, known as IVS-110, generates an *HBB* mRNA with an in-frame stop codon, resulting in a 90% reduction in functional HBB protein through nonsense-mediated decay (Spritz et al., 1981; Vadolas et al., 2006). We utilized two MEL cell lines expressing either an integrated copy of a human β-globin minigene (MEL-*HBB*^WT^) or the human β-globin minigene with the IVS-110 mutation (MEL-*HBB*^IVS-110(G>A)^) (Patsali et al., 2018). Specific targeting of these integrated human *HBB* loci during library preparation resulted in an average of 24,970 nascent RNA long-reads that mapped to the *HBB* gene for each of 3 biological replicates (**Table S1**), allowing rigorous statistical analysis. As previously reported, the majority (94%) of intron 1 splicing in the MEL-*HBB*^IVS-110(G>A)^ cell line occurred at the cryptic 3’SS.

**Figure 7.**
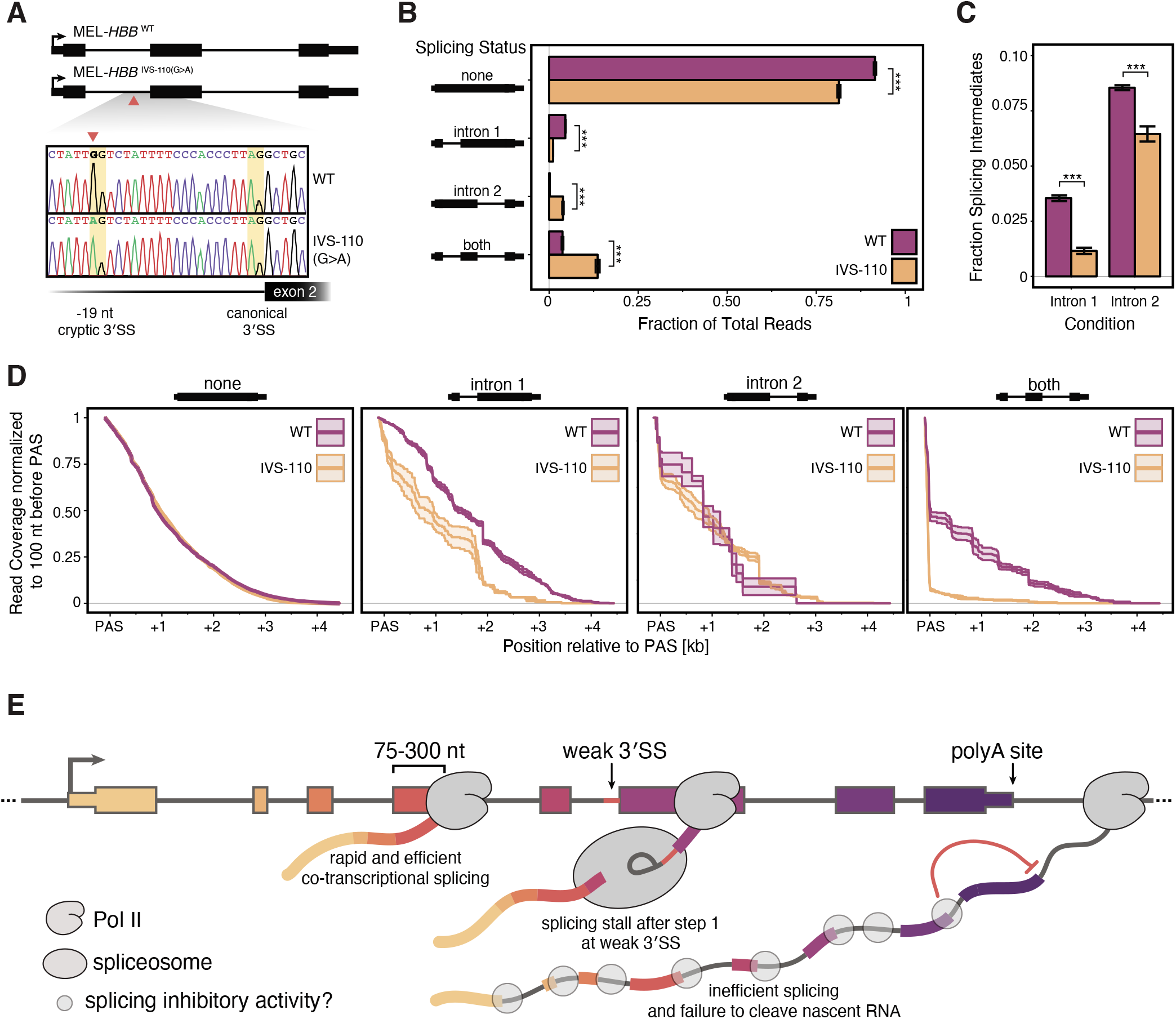
Efficient splicing promotes 3’ end cleavage. **(A)** Top: schematic describing two engineered MEL cell lines. MEL-*HBB* ^WT^ contains an integrated copy of a wild type human globin minigene. In MEL-*HBB*^IVS-110(G>A)^, a single point mutation (red triangle) mimics a disease-causing thalassemia allele. Bottom: Sanger sequencing of the *HBB* minigene coding strand shows that a G>A mutation leads to a cryptic 3’SS at the AG dinucleotide 19 nt upstream of the canonical 3’SS. **(B)** Distribution of *HBB* long-reads in MEL-*HBB* ^WT^ cells (purple) and MEL-*HBB*^IVS-110(G>A)^ cells (orange) separated by splicing status of intron 1 and intron 2 and measured as a fraction of total reads mapped to the *HBB* gene (n = 20,395 reads in MEL-*HBB* ^WT^ cells, and n = 26,244 reads in MEL-*HBB*^IVS-110(G>A)^ cells). **(C)** Fraction of splicing intermediates at intron 1 and intron 2 in MEL-*HBB* ^WT^ cells (purple) and MEL-*HBB* ^IVS-110(G>A)^ cells (orange) measured as a fraction of total reads mapped to the *HBB* gene. For **(B-C)**, significance tested by Mann Whitney U-test; *** represents p-value < 0.001, bar height represents the mean of three biological replicates, and error bars represent standard error of the mean. **(D)** Read coverage in the region downstream of the *HBB* PAS is shown for long-reads separated by their splicing status from MEL-*HBB* ^WT^ cells (purple) and MEL-*HBB* ^IVS-110(G>A)^ cells (orange). Coverage is normalized to the position 100 nt upstream of the PAS. Solid line represents the mean coverage of three biological replicates, and shaded windows represent standard error of the mean. **(E)** Model describing the variety of co-transcriptional splicing efficiencies observed during mouse erythropoiesis.

*MEL-HBB*^IVS-110(G>A)^ cells exhibited a significant increase in the fraction of long-reads that were all spliced and a corresponding decrease in long-reads that were all unspliced (**Figure 7B**). The low co-transcriptional splicing efficiency detected for endogenous mouse β-globin was mirrored in the stably integrated *HBB* minigene, where 80-92% of long-reads were all unspliced (compare **Figures 6A and 7B**). Long-reads with only intron 1 spliced were present at a higher ratio in the MEL-*HBB* ^WT^ cells, whereas long-reads with only intron 2 spliced were more frequent in the MEL-*HBB*^IVS-110(G>A)^ cells. This distribution suggests that the cryptic intron 1 is spliced more efficiently than the WT intron 1, leading to a coordinated increase in splicing of intron 2 and a shift from all unspliced to all spliced reads. This interpretation agrees with the coordinated co-transcriptional splicing efficiency we observed in multi-intron transcripts genome-wide (**Figure 2E**). Significantly fewer splicing intermediates were detected for both introns in the MEL-*HBB*^IVS-110(G>A)^ cells compared to MEL-*HBB* ^WT^ cells, suggesting that inefficient splicing can lead to spliceosomal pausing between catalytic steps I and II (**Figure 7C**).

To rigorously test the possibility that changes in co-transcriptional splicing efficiency determine 3’ end cleavage, read coverage downstream of the *HBB* PAS was used to detect uncleaved long-reads for each category of splicing status (**Figure 7D**). All-unspliced *HBB* reads were detected up to 4 kb past the PAS, similar to endogenous mouse globin genes. When only intron 2 was spliced, cleavage in MEL-*HBB* ^WT^ and MEL-*HBB*^IVS-110(G>A)^ cells was similar (**Figure 7D**; center right). However, a notable decrease in uncleaved reads past the PAS was detected among transcripts spliced at the cryptic 3’SS as compared to the canonical 3’SS (**Figure 7D**; center left). When both introns were spliced, there was an even more dramatic shift to proper cleavage at the PAS in the MEL-*HBB*^IVS-110(G>A)^ cells (**Figure 7D**; far right). Together, these findings confirm our demonstration of functional coupling between splicing and 3’ end formation at the individual transcript level and highlight the regulatory potential of just a single point mutation. Thus, a previously unappreciated level of crosstalk between splicing and 3’ end cleavage efficiencies is involved in erythroid development.

### DISCUSSION

This study reveals functional relationships between co-transcriptional RNA processing events through genome-wide analysis of individual nascent transcripts purified from differentiating mammalian erythroid cells. Transcription and splicing dynamics were visualized with unprecedented depth and accuracy through long-read sequencing of nascent RNA and PRO-seq. We conclude that splicing catalysis can occur when Pol II is just 75-300 nt past the intron without transcriptional pausing at the splice sites. Thus, spliceosome assembly and the transition to catalysis often occur when the spliceosome is physically close to Pol II. Two striking cases stood out from our observations of splicing. First, introns that contain a weak 3’SS seem to induce stalling between the steps I and II of the splicing reaction itself, causing a buildup of splicing intermediates. Second, inefficient splicing was globally correlated with inefficient 3’ end cleavage on both the population and single transcript level (**Figure 7E**). We pursued this second phenomenon further in the context of globin gene expression, wherein all two-intron globin genes (two α and two β in mouse) displayed “all-or-none” splicing behavior. Approximately 20% of endogenous nascent *Hbb-b1* transcripts retained both introns and were inefficiently cleaved at the PAS. Remarkably, a patient-derived, thalassemia-causing point mutation in β-globin increased splicing efficiency and 3’ end cleavage. These data show that co-transcriptional splicing efficiency determines 3’ end processing efficiency, as discussed below.

The data presented indicate that the mammalian spliceosome is capable of assembling and acting on nascent RNA substrates in the same spatial window of transcription as the yeast spliceosome (Carrillo Oesterreich et al., 2016; Herzel et al., 2018). This is despite changes in the complexity of yeast and mammalian spliceosomes, the length and number of introns in mammals, as well as the number of accessory factors that can influence splicing. Our high resolution determination of the fraction of splicing that occurs co-transcriptionally (88%) matches data from alternative methods (e.g. short-read sequencing, metabolic labeling, and imaging) showing that 75-87% of splicing is co-transcriptional in yeast, fly, mouse, and human cells (reviewed in (Neugebauer, 2019). These findings suggest widely conserved features of transcription and splicing mechanisms.

A recent paper reports that the majority of introns in both human and fly nascent RNA appear unspliced and that Pol II is 2-4 kb downstream of the 3’SS when splicing occurs (Drexler et al., 2020), which is at odds with our findings. One possible explanation for this discrepancy is that the purification of 4sU-labeled RNA inadvertently enriched for long, intron-containing RNAs that have a greater probability of containing a labeled U residue (introns are U-rich). 4sU incorporation may also impede splicing due to changes in base-pairing among U-rich RNA elements in introns and snRNAs (Testa et al., 1999). Indeed, direct comparison of splicing efficiencies with vs. without 4sU-labelling indicates that 4sU-lableled RNA is less frequently spliced (Drexler et al., 2020). In this context, we note that the principal difference between the data from our two labs is the fraction of unspliced RNAs observed. However, when we analyze spliced RNAs in the Drexler *et al.* data to determine the distance between splice junctions and the RNA 3’ end, we obtained similar results to our own (**Figure S4A**). We conclude that splicing more typically occurs when Pol is close to the intron.

Importantly, PRO-seq corroborated the efficiency of co-transcriptional splicing. We identified spliced reads within the PRO-seq data, validating the observations made with LRS of purified nascent RNA with an independent method. Similarly, mNET-seq data, which is generated by short-read sequencing of nascent RNA from immunoprecipitated Pol II, has revealed examples of spliced reads (Nojima et al., 2018). Having observed many examples wherein an RNA 3’ end was only a short distance beyond the 3’SS, we considered the hypothesis that Pol II pausing at or near 3’SSs could provide extra time for splicing (Alexander et al., 2010b; Chathoth et al., 2014; Milligan et al., 2017). However, our analysis shows that any detection of a PRO-seq peak at 5’SSs—albeit small in meta-analysis—is caused by bleed-through from promoter-proximal pausing, and we do not detect statistically significant pausing at 5’ or 3’SSs (**Figure 4A&B**), in agreement with a recent study using mNET-seq (Sheridan et al., 2019). Importantly, Pol II elongation was not detectably impacted by the splicing efficiency of introns (**Figure S5C**), with no significant differences in PRO-seq signal across splice junctions. These data strongly support the assumption that Pol II travels at a uniform rate across splice junctions. Using the median distance from splicing events to Pol II position in our combined data (142 nt) and taking into account the 0.5-6 kb/min range of measured Pol II elongation rates (Jonkers and Lis, 2015), we calculate that the splicing events detected in our data occurred within 1.4-17 seconds on intron transcription. These rates are similar to those obtained in budding and fission yeasts (Alexander et al., 2010a; Alpert et al., 2020; Carrillo Oesterreich et al., 2016; Eser et al., 2016; Herzel et al., 2018).

Due to the 3’ end chemistry and structure of splicing intermediates, we can capture the step I intermediates of splicing with long-read sequencing. Remarkably, splicing intermediates were distributed unevenly among introns and were associated with poor sequence consensus at the downstream 3’SS. This evidence is consistent with a model where modulation of the transition between catalytic steps of splicing can alter splicing fidelity or outcome (Smith et al., 2008). Because the spliceosomes associated with these intermediates have already assembled and undergone step 1 chemistry, the accumulation of intermediates cannot be attributed to defective intron recognition during spliceosome assembly. Instead, the catalytic center of the spliceosome shifts from the branch site (step I) to the 3’SS AG (step II), typically 30-60 nt downstream of the branch site. The mechanism underlying 3’SS choice during this transition is likely to be influenced by spliceosomal proteins outside of the catalytic core. A recent cryo-EM study of human spliceosomes has identified several spliceosomal components that may be in a position to regulate the transition from step I to step II (Fica et al., 2019). Future studies of these enigmatic new players may reveal a role for 3’SS diversity in the regulation of splicing by stalling between catalytic steps.

Our LRS data resolve a mystery shrouding □-globin pre-mRNA splicing. Two previous studies used stably integrated □-globin reporter genes combined with high resolution fluorescence microscopy to track pre-mRNA transcription and splicing in HEK293 and U2OS cells (Coulon et al., 2014; Martin et al., 2013). One study reported data consistent with co-transcriptional splicing, while the other strongly favored post-transcriptional splicing. The LRS data presented here explains that, at least in MEL cells, there are two major populations of globin transcripts: “all spliced” RNAs and “all unspliced” RNAs (**Figure 6A; Figure 7B; Figure S7A**). Although previous studies also linked the splicing of □-globin exon 2 with 3’ end cleavage (Antoniou et al., 1998; Dye and Proudfoot, 1999), here we show all-or-none behavior in the splicing and cleavage decisions made by both □- and □-globins. In any biochemical and/or short-read RNA-seq assay that examines populations of pre-mRNA, inefficient splicing would be one explanation for the bulk result. In reality, one population is unspliced and uncleaved at their 3’ ends. The other population of globin transcripts is efficiently spliced and productively expressed, because polyA cleavage and subsequent Pol II termination also occur efficiently for these RNAs.

The fraction of efficiently spliced □-globin transcripts increased in the thalassemia allele we studied, even though the cryptic 3’SS yields an out of frame mRNA that will - like many thalassemia alleles of □-globin - be degraded by nonsense-mediated decay (Kurosaki et al., 2019). Accordingly, a decrease in uncleaved reads past the PAS in the MEL-*HBB*^IVS-110(G>A)^ cells was evident compared to the MEL-*HBB* ^WT^ cells. This indicates that splicing efficiency is a determinant of 3’ end cleavage. Several possible mechanisms could be involved. Less efficient splicing can inhibit 3’ end cleavage (Cooke et al., 1999; Davidson and West, 2013; Martins et al., 2011), suggesting that introns retained in transcripts that display readthrough harbor an inhibitory activity that represses 3’ end cleavage (**Figure 7E**). Candidate inhibitory factors include U1 snRNP, PTB and hnRNP C proteins, each of which binds introns promiscuously *in vivo* (Deng et al., 2020; König et al., 2010). In particular, U1 snRNP binding to introns is known to repress premature 3’ end cleavage and cleavage at PASs (Berg et al., 2012; So et al., 2019; Vagner et al., 2000). We speculate that this inhibitory activity persists longer on inefficiently spliced transcripts, potentially binding and inactivating 3’ end cleavage factors (Deng et al., 2020; So et al., 2019). Additionally, the deposition of stimulatory factors - namely the exon junction complex (EJC) and SR proteins - on exon-exon junctions after splicing may stimulate splicing of the next intron (Singh et al., 2012), potentially contributing to the all-spliced phenomenon. An added stimulus to 3’ end cleavage may also be afforded by loss of U1 snRNP and accumulation of SR proteins, since the RS domain in Fip1 promotes cleavage (Zhu et al., 2018); more generally, SR proteins are associated with polyA site choice (Muller-McNicoll et al., 2016). Thus, more efficient splicing in the thalassemia mutant likely enables 3’ end cleavage by more quickly removing inhibitory activities and/or recruiting positive effectors. Investigation of these mechanisms awaits future studies that would afford single transcript evaluation of the residence time of intron-bound inhibitory factors (e.g. U1 snRNP) coupled with splicing and cleavage outcome.

It is tempting to speculate that improving splicing efficiency could be a general strategy for increasing gene output in a variety of disease settings. Our findings substantiate an earlier proposal based on experiments on □-globin that efficient splicing and 3’ end cleavage contribute to gene expression output (Lu and Cullen, 2003), by suggesting that nascent transcripts are earmarked as productive or unproductive during their biogenesis. Moreover, failure of 3’ end cleavage when splicing is impaired would explain why splicing efficiency was previously associated with release of RNA from the site of transcription (Antoniou et al., 1998; Custodio et al., 1999; Dye and Proudfoot, 1999). Importantly, many physiological stresses – such as osmotic stress, heat shock, cancer, aging, and viral infection – cause a failure to cleave at annotated PASs and the production of very long non-coding RNAs (Enge et al., 2017; Grosso et al., 2015; Muniz et al., 2017; Vilborg et al., 2015; Vilborg et al., 2017). This strong connection between splicing efficiency, 3’ end formation, and transcription termination introduces previously unknown layers of regulation to mammalian gene expression in a variety of physiological contexts.

### LIMITATIONS

Several limitations to this study remain to be investigated further. First, the length of long-reads are dependent on reverse transcriptase processivity when copying RNA into cDNA. While we have taken steps to enrich for full-length transcripts in our library generation, some RNAs are likely not fully reverse transcribed and captured in this dataset. Advancements in strand-switching RT enzyme chemistry may improve this in the future (Guo et al., 2020). Second, we have not addressed directly what the ultimate fate of unspliced and uncleaved nascent RNA is in these cells. While in other studies, we found these transcripts were degraded by the nuclear exosome (Herzel et al., 2018), it remains to be tested directly. Finally, a more rigorous test of our proposed mechanism linking splicing and 3’ end cleavage would require tools to probe inhibition of both processes. While chemical inhibitors can be used to block spliceosome assembly globally, these drugs also induce a general stress response in cells (Castillo-Guzman et al., 2020), including changes in transcription and 3’ end cleavage. Thus, future studies probing this mechanism await specific reagents to test directly the link between splicing and cleavage.

## ACKNOWLEDGMENTS

We thank P Patsali for sharing the MEL-*HBB* ^WT^ and MEL-*HBB*^IVS-110(G>A)^ cell lines, M Antoniou for sharing an annotation of the GLOBE vector, and J Conboy for advice on erythroblast fractionation. We thank E Brown for help with preparation of LRS figures, J Gordon for technical assistance, and H Tilgner, T Carrocci, D Phizicky, T Alpert, T Henriques, and B Martin for helpful discussions and comments on the manuscript. This work was initiated through pilot funding from NIDDK under Grant U54DK106857 to the Yale Cooperative Center of Excellence in Hematology (to K.M.N.). It was further supported by the National Institutes of Health (NIH R01 GM112766 to K.M.N) and Startup Funding from Harvard Medical School (to K.A). Its contents are solely the responsibility of the authors and do not necessarily represent the official views of the NIH. K.A.R. is supported by a Postgraduate Scholarship from the Natural Sciences and Engineering Research Council of Canada (NSERC) and a Gruber Science Fellowship, and C.M. is supported by a National Science Foundation Graduate Research Fellowship (DGE1745303).

## AUTHOR CONTRIBUTIONS

Conceptualization, K.M.N. and K.A.R.; Investigation, K.A.R. and C.M.; Data Curation, K.A.R. and C.M.; Writing -- Original Draft, K.A.R. and K.M.N.; Writing -- Review & Editing, K.A.R., C.M., K.A., and K.M.N.; Visualization, K.A.R. and C.M.; Supervision, K.A. and K.M.N.; Funding Acquisition, K.A. and K.M.N.

---

**Figure S1.**
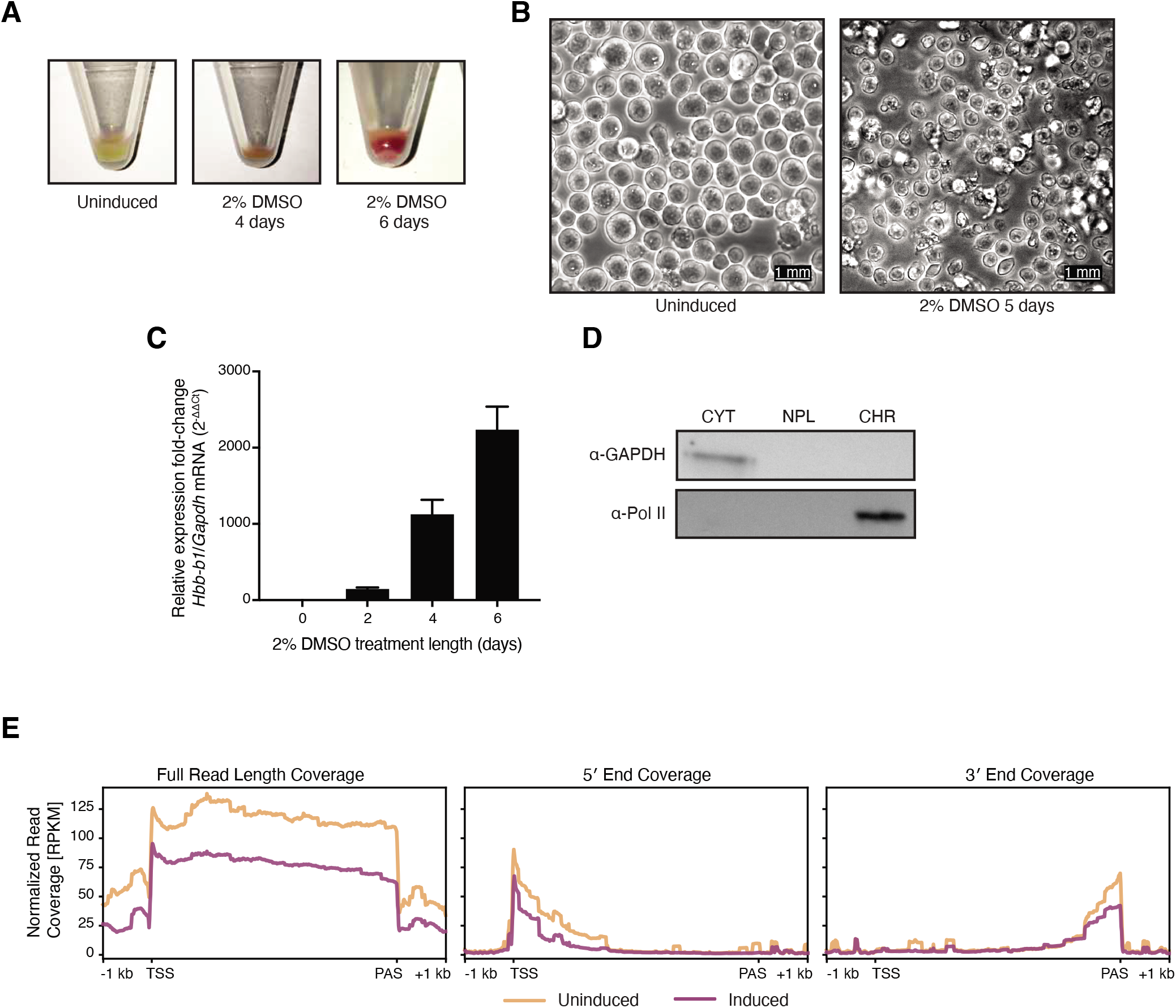
Related to Figure 1. DMSO treatment induces erythroid differentiation. **(A)** MEL cells after culturing in 2% DMSO for 0, 4, or 6 days. **(B)** Bright field microscopy of MEL cells uninduced (left) and induced for 5 days (right). Scale bar is 1 mm. **(C)** RT-qPCR measurement of *Hbb-b1* (β-globin) mRNA levels relative to *Gapdh* mRNA from total RNA in MEL cells treated with 2% DMSO for 0, 2, 4, or 6 days. Bar heights represent mean of 3 technical replicates, and error bars represent SEM. **(D)** Western blot of subcellular fractions collected during chromatin fractionation (CYT = cytoplasm, NPL = nucleoplasm, CHR = chromatin). **(E)** Normalized read coverage (RPKM) of long-read 5’ ends (left), full reads (middle), and 3’ ends (right) is shown across a metaplot of all mm10 genes +/− 1 kb (TSS = transcription start site, TES = transcription end site).

**Figure S2.**
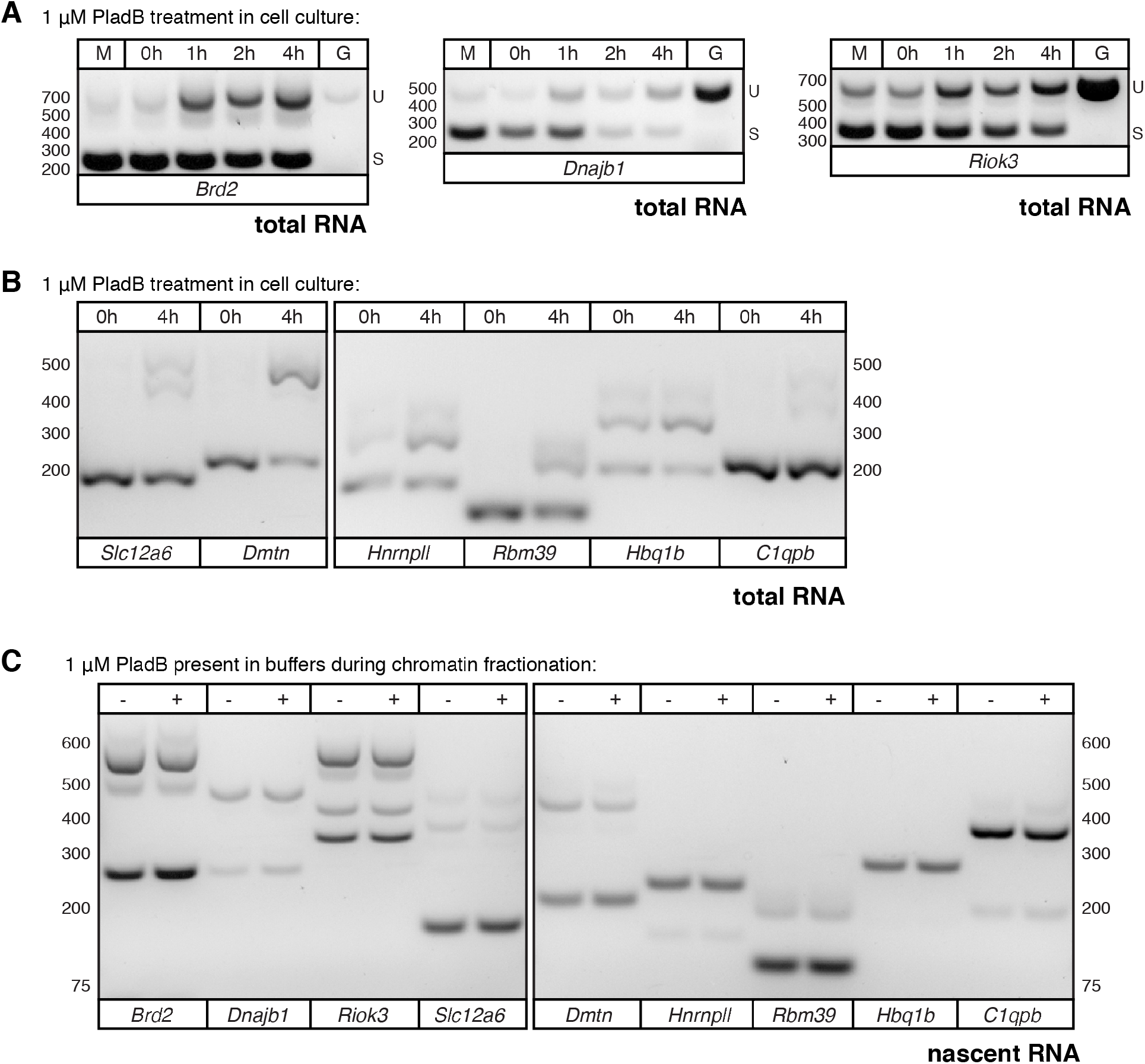
Related to Figure 1. Splicing does not continue during chromatin purification and nascent RNA isolation. **(A)** RT-PCR on total RNA collected from MEL cells treated with 1 μM splicing inhibitor Pladienolide B in cell culture for 0, 1, 2, and 4 hours. Total RNA was reverse transcribed with random hexamers, and PCR primers span a single intron in each gene. Three representative genes are shown (left: *Brd2,* middle: *Dnajb1,* and right: *Riok3*). M indicates mock treatment with DMSO, and G indicates amplification of genomic DNA to determine the size of unspliced RNA. U indicates size of unspliced amplicon, and S indicates size of spliced amplicon. **(B)** RT-qPCR from total RNA (as in **(A)**), showing six additional genes (*Slc12a6, Dmtn, Hnrnpll, Rbm39, Hbq1b, C1qbp*) after treatment with 1 μM Pladienolide B for 0h and 4h. **(C)** RT-PCR on nascent RNA isolated from chromatin which was fractionated in the absence (-) or presence (+) of 1 μM Pladienolide B. Nascent RNA was reverse transcribed with random hexamers and PCR primers were the same as in **(A)** and **(B)**.

**Figure S3.**
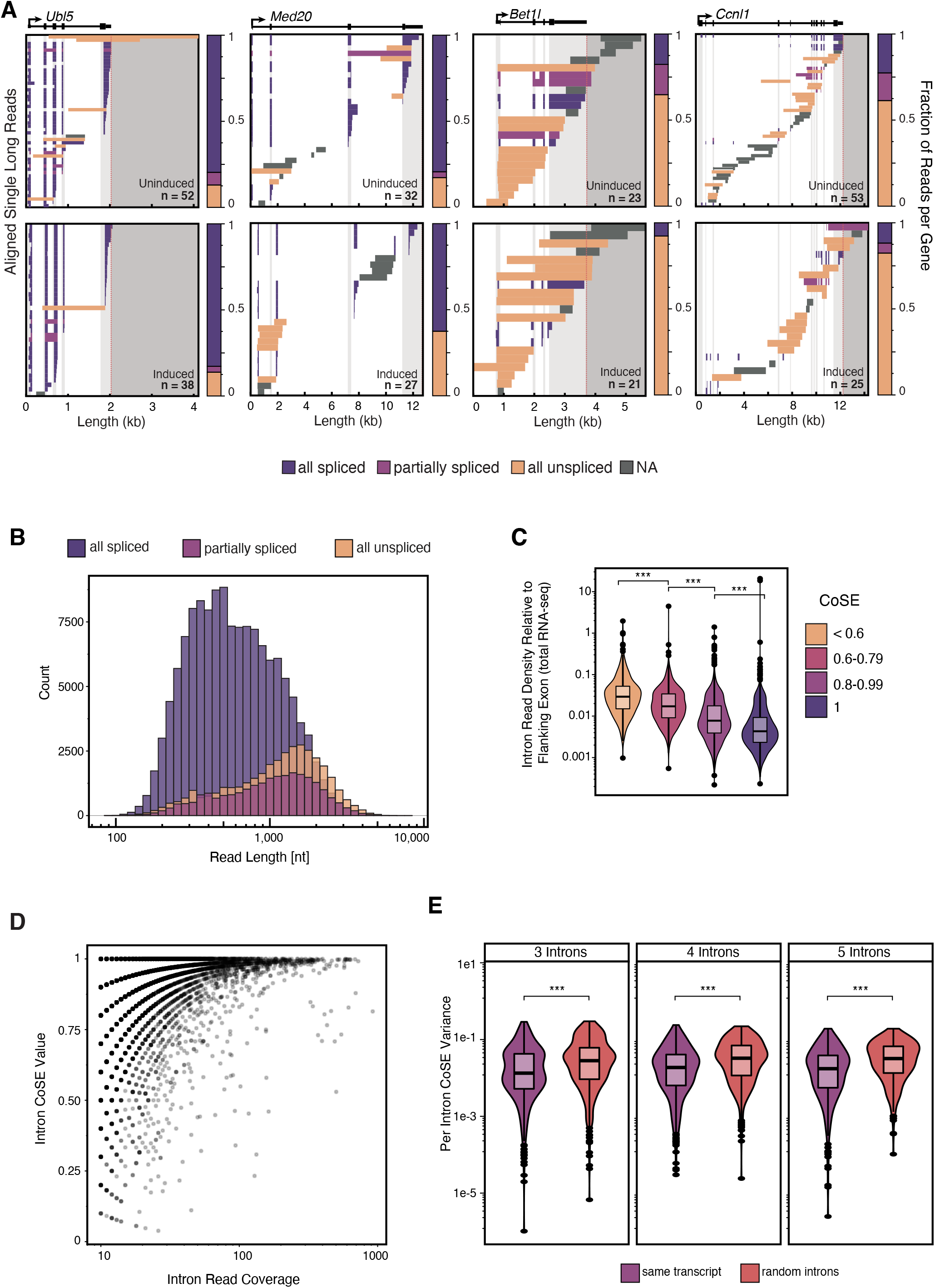
Related to Figure 2. Long-read sequencing reveals similar co-transcriptional splicing efficiencies (CoSE) for introns within the same transcript. **(A)** Individual long-reads for genes *Ub15, Med20, Bet1l, and Ccnl1* are shown as described in **Figure 2**. **(B)** Long-read length distribution separated by splicing status (n = 134,857 all spliced reads, n = 23,833 partially spliced reads, n = 35,957 all unspliced reads). **(C)** Intron coverage relative to flanking exon coverage from untreated MEL total RNA-seq Illumina data (ENCODE) is shown as an independent indicator of unspliced RNA levels. Introns that were covered by at least 10 long-reads in the untreated condition were split into bins based on the CoSE metric calculated from long-read sequencing data. Significance tested by Mann Whitney U-test; *** represents p-value <0.001. **(D)** CoSE values as a function the number of long-reads covering each intron. Each point represents an intron for which a CoSE value was calculated, with uninduced and induced data shown combined (n = 14,422 introns). **(E)** Variance in CoSE for transcripts that include 3, 4, or 5 introns with at least 10 reads (“same transcript”) compared to the variance in CoSE for a randomly selected group of introns. Randomly chosen introns were grouped such that the groupings match the number of introns per transcript in the “same transcript” data (3 introns n = 455, 4 introns n = 363, 5 introns n = 266). For **C** and **E**, significance tested by Mann Whitney U-test; *** represents p-value < 0.001.

**Figure S4.**
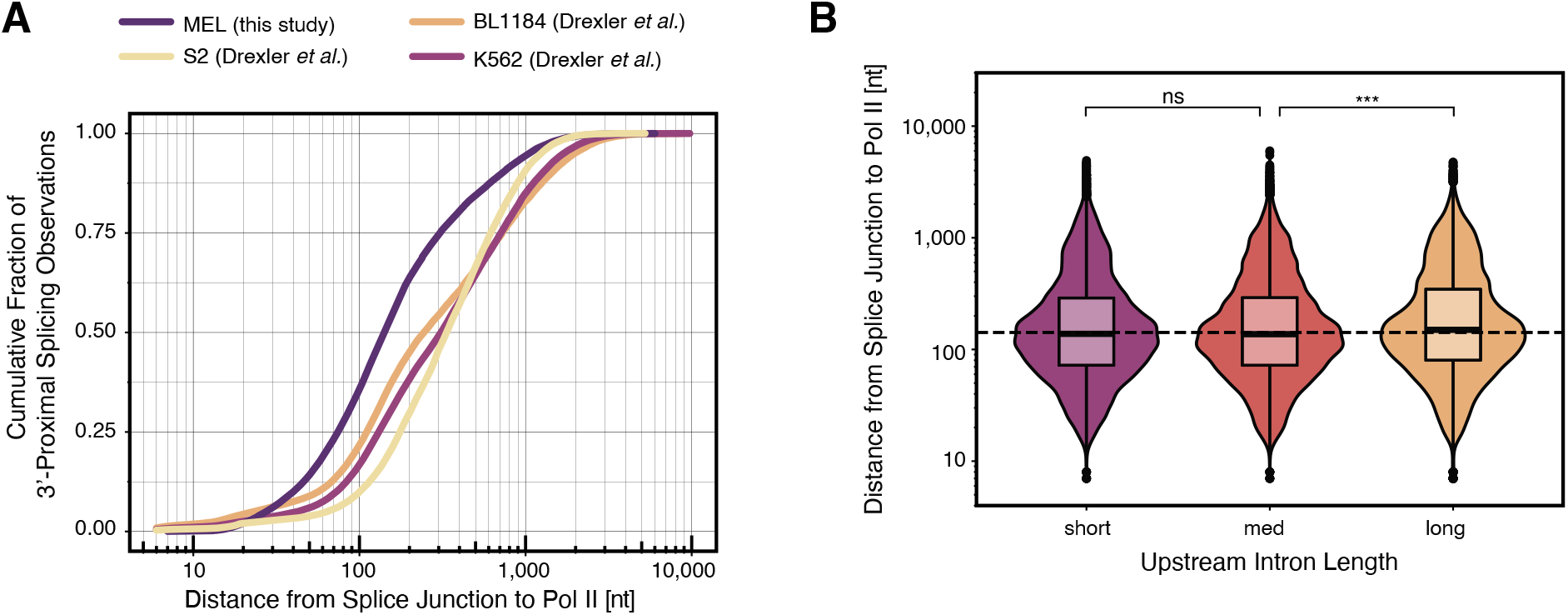
Related to Figure 3. Pol II is detected near splice junctions in multiple cell types. **(A)** Distance (nt) from the most 3’-proximal splice junction to Pol II position is shown as a cumulative fraction. Analysis is the same as in **Figure 3B**, but with data from this study and nanoCOP data from (Drexler et al., 2020) (n = 39,397 reads in BL1184 cells, n = 183,829 reads in K562 cells, n = 68,956 reads in S2 cells, and n = 168,567 reads in MEL cells). **(B)** Distance (nt) from the most 3’-proximal splice junction to Pol II position is shown categorized by upstream intron length in three equal-sized bins (short = 30 nt - 443 nt, med = 443 nt - 1,738 nt, long = >1,738 nt). Significance tested by Mann Whitney U-test: *** represents p-value < 0.001, ns represents p-value > 0.05.

**Figure S5.**
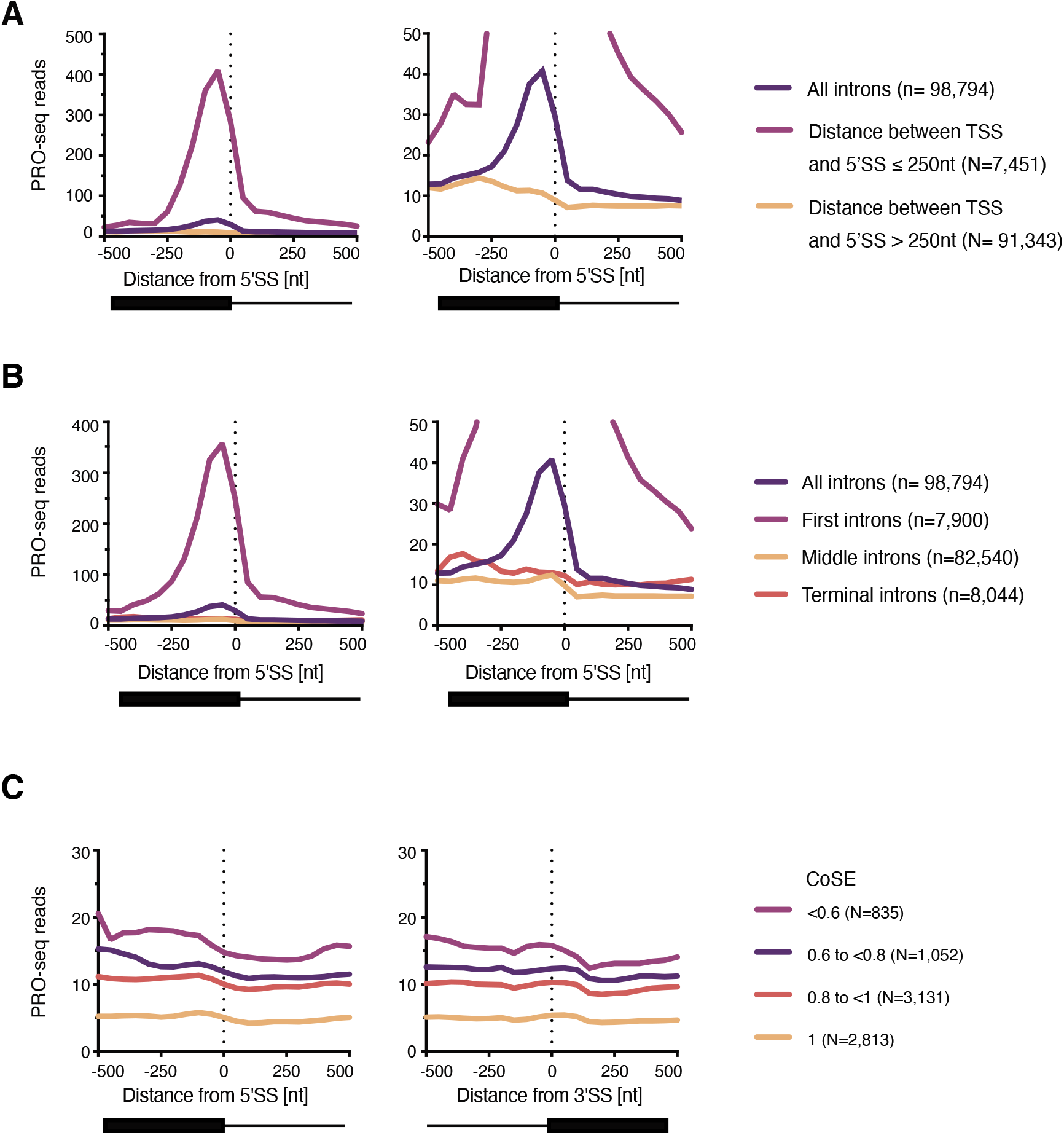
Related to Figure 4. PRO-seq reveals no Pol II pause at any subset of introns. **(A)** PRO-seq 3’ end coverage aligned to 5’SSs for all introns (dark purple), introns where the distance from the TSS to the 5’SS is ≤ 250 nt (light purple), and introns where the distance to the TSS to the 5’SS is > 250 nt (light orange). Right panel shows data scaled to show all introns. **(B)** PRO-seq 3’ end coverage aligned to 5’SSs for all introns from active transcripts (dark purple), first introns only (light purple), middle introns (light orange), and terminal intron (dark orange). Right panel shows data scaled to show all introns. **(F)** Average PRO-seq 3’ end coverage is shown around 5’SS (left) and 3’SS (right) for introns in each CoSE category as indicated. N = number of introns in each category.

**Figure S6.**
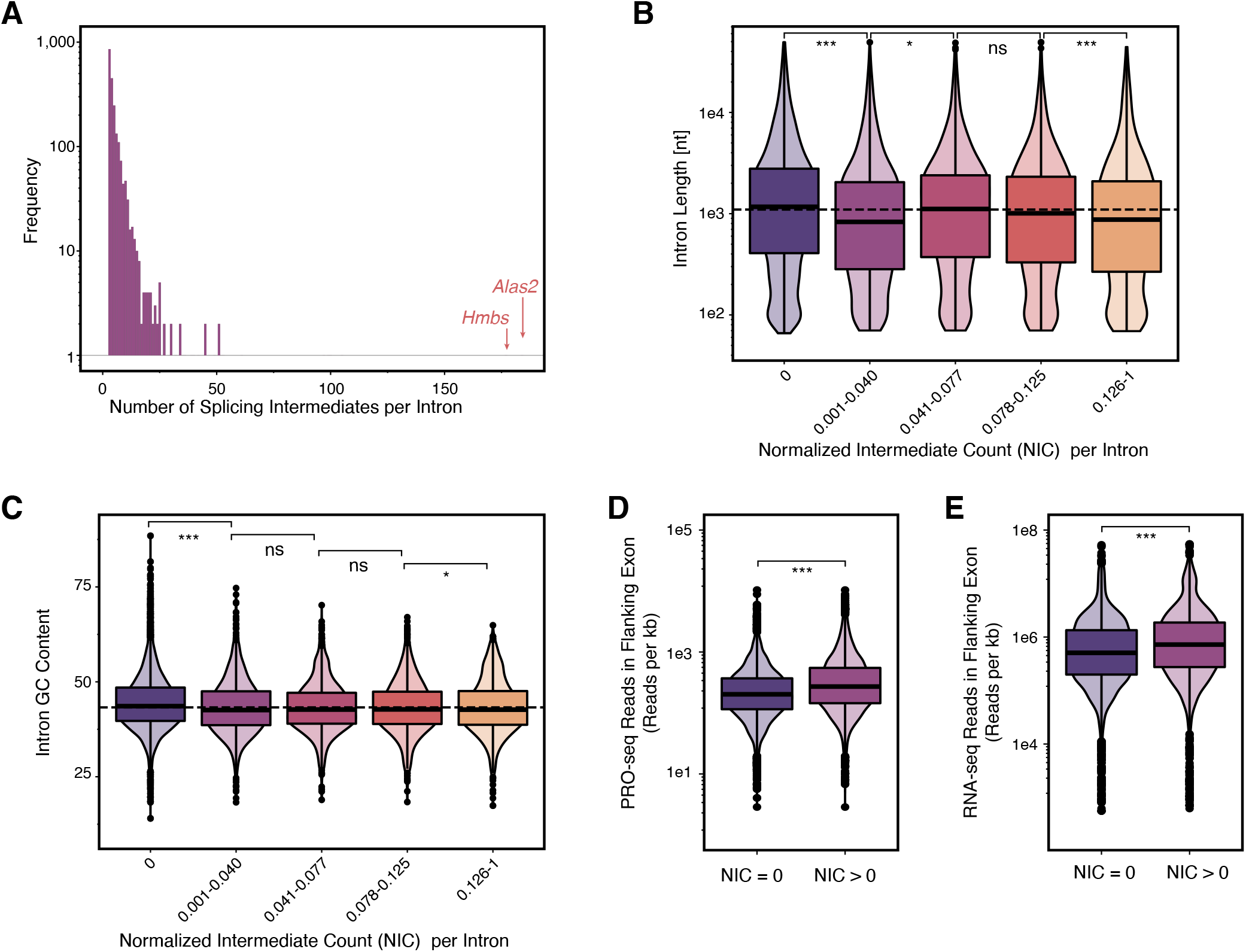
Related to Figure 5. Splicing intermediates are rare at most introns, but highly abundant at a few introns. **(A)** Histogram showing frequency of splicing intermediates upstream of each unique intron from active transcripts in MEL cells. Most introns show 0 or 1 splicing intermediates, while some introns, like those in *Alas2* and *Hmbs,* indicated with orange arrows, exhibit over 150 splicing intermediates reads. **(B)** Intron length (nt) and **(C)** intron GC-content for unique introns from active transcripts categorized by NIC value. I Introns with NIC = 0 (n = 3,890) are shown separately, and all other introns with NIC > 0 (n = 2,647) are separated in quartiles with NIC values shown. **(D)** PRO-seq and **(E)** Total RNA-seq (ENCODE) reads summed within the upstream and downstream exons surrounding each intron containing 10 or more long-reads in the uninduced condition and normalized by exon length for introns with NIC = 0 (dark purple) or NIC > 0 (light purple). For **B-E**, significance tested by Mann Whitney U-test; *** represents p-value < 0.001, ns represents p-value > 0.05. For **(A-C)**, data represent two biological and two technical replicates from induced and uninduced cells combined.

**Figure S7.**
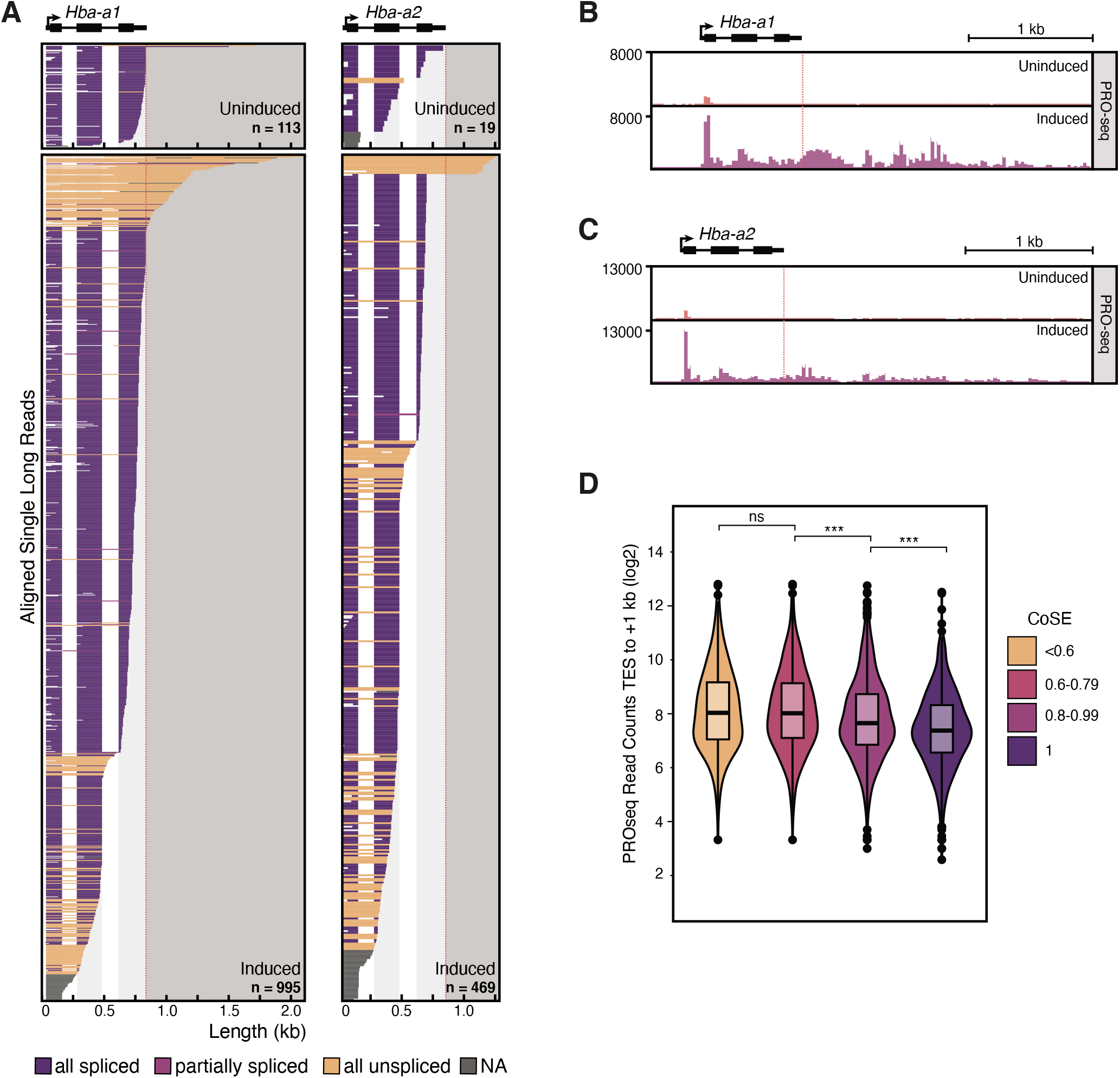
Related to Figure 6. α-globin genes exhibit unspliced transcripts that are uncleaved and extend past the PAS. **(A)** Individual long-reads are shown for the α-globin 1 gene (*Hba-a1*) and α-globin 2 gene (*Hba-a2*). Diagrams are as described in **Figure 2**. Data represent two biological replicates and two technical replicates combined. **(B-C)** PRO-seq 3’ end read coverage is shown downstream of the *Hba-a1* **(B)** and *Hba-a2* **(C)** gene loci. Note that the duplicated copies of α-globin in the genome (*Hba-a1* and *Hba-a2*) impedes unique mapping of short PRO-seq reads in the coding sequence, artificially reducing the density of gene body reads. Red dotted line indicates PAS. Data represent three biological replicates combined. **(D)** Summed PRO-seq signal in the region 1 kb downstream of the PAS in uninduced cells. Introns were binned by CoSE values, then the PRO-seq read coverage was calculated for each intron and counted only once per bin. Significance tested by Mann-Whitney U-test; *** represents p-value < 0.001, ns represents p-value > 0.05.

## STAR METHODS

### RESOURCE AVAILABILITY

#### Lead Contact

Further information and requests for resources and reagents should be directed to and will be fulfilled by the Lead Contact, Karla Neugebauer.

#### Materials Availability

This study did not generate new unique reagents.

#### Data and Code Availability

Raw and processed long-read sequencing and PRO-seq data generated in this study are deposited in NCBI’s Gene Expression Omnibus and are accessible through GEO Series accession number **GSE144205**. Raw image data associated with this manuscript are available on Mendeley (http://dx.doi.org/10.17632/5vrtbpnj4k.1). All code supporting the long-read sequencing data analysis in this manuscript is available at https://github.com/NeugebauerLab/MEL_LRS. NanoCOP data from Drexler *et al.* 2020 analyzed in this manuscript can be found at GEO with accession number **GSE123191**, and total RNA-seq from MEL cells analyzed in this study can be found at Mouse ENCODE (http://www.mouseencode.org/) with accession number **ENCSR000CWE**.

### EXPERIMENTAL MODEL AND SUBJECT DETAILS

#### Cell lines, cell culture, and treatments

Murine Erythroleukemia cells (MEL; obtained from Shilpa Hattangadi, Yale School of Medicine) were maintained at 37°C and 5% CO_2_ in DMEM + Glutamax medium (GIBCO) containing 100 U/ml penicillin, 100 μg/ml streptomycin (GIBCO), and 10% fetal bovine serum (GIBCO). To induce erythroid differentiation, cells were diluted to 50,000 cells/ml in 10 ml fresh culture medium and incubated as above for 16 hours. DMSO was then added directly to the culture medium to a final concentration of 2% and incubated as above for 5 days. For Pladienolide B treatment, cells were diluted to 50,000 cells/ml in fresh culture medium, then incubated as above for two days until reaching a density of approximately 5 million cells/ml. Pladienolide B (Santa Cruz) dissolved in DMSO was added directly to the culture medium at a final concentration of 1 μM. MEL-*HBB* ^WT^ and MEL-*HBB*^IVS-110(G>A)^ cell lines are described previously (Patsali et al., 2018), and were maintained and differentiated as above.

### METHOD DETAILS

#### Subcellular Fractionation

Subcellular fractionation was adapted from previously published protocols (Mayer and Churchman, 2017; Pandya-Jones and Black, 2009), with modifications to centrifugation speeds in order to retain intact nuclei (Reimer and Neugebauer, 2020). All steps were performed on ice, and all buffers contained 25 uM α-amanitin, 40 U/ml SUPERase.IN, and 1x Roche cOmplete protease inhibitor mix. Briefly, 20 million cells were rinsed once with PBS/1 mM EDTA, then lysed in 250 μl cytoplasmic lysis buffer (10 mM Tris-HCl pH 7.5, 0.05% NP40, 150 mM NaCl) by gently resuspending then incubating on ice for 5 minutes. Lysate was then layered on top of a 500 μl cushion of 24% sucrose in cytoplasmic lysis buffer and spun at 2000 rpm for 10 min at 4°C. The supernatant (cytoplasm fraction) was removed, and the pellet (nuclei) were rinsed once with 500 μl PBS/1 mM EDTA. Nuclei were resuspended in 100 μl nuclear resuspension buffer (20 mM Tris-HCl pH 8.0, 75 mM NaCl, 0.5 mM EDTA, 0.85 mM DTT, 50% glycerol) by gentle flicking, then lysed by the addition of 100 μl nuclear lysis buffer (20 mM HEPES pH 7.5, 1 mM DTT, 7.5 mM MgCl_2_, 0.2 mM EDTA, 0.3 M NaCl, 1 M Urea, 1 % NP-40), vortexed for 2 x 2 seconds, then incubated on ice for 3 min. Chromatin was pelleted by spinning at 14,000 rpm for 2 min at 4°C. The supernatant (nucleoplasm fraction) was removed, and the chromatin was rinsed once with PBS/1 mM EDTA. Chromatin was immediately dissolved in 100 μl PBS and 300 μl TRIzol Reagent (ThermoFisher).

#### Nascent RNA Isolation

RNA was purified from chromatin pellets in TRIzol Reagent (ThermoFisher) using the RNeasy Mini kit (Qiagen) according to the manufacturer’s protocol, including the on-column DNase I digestion. For genome-wide nascent RNA-seq, samples were depleted three times of polyA(+) RNA using the Dynabeads mRNA DIRECT Micro Purification Kit (ThermoFisher), each time keeping the supernatant, then depleted of ribosomal RNA using the Ribo-Zero Gold rRNA Removal Kit (Illumina). For targeted nascent RNA-seq, polyA(+) and rRNA depletion were omitted.

#### Western Blotting

Cytoplasm, nucleoplasm, and chromatin fractions from cell fractionation were adjusted to an equal volume with PBS. Nucleoplasm and chromatin fractions were homogenized by sonication, and all samples were spun at 14,000 rpm for 10 min at 4°C before gel loading. Western blots were performed with antibodies against Pol II 4H8 (Santa Cruz Biotechnology) and GAPDH (Santa Cruz Biotechnology).

#### qPCR

Total RNA was extracted from 10 million cells in TRIzol Reagent (ThermoFisher) according to the manufacturer’s protocol after 0, 2, 4, or 6 days of treatment with 2% DMSO as described above. cDNA was generated with SuperScript III Reverse Transcriptase (ThermoFisher) using random hexamer primers (ThermoFisher) according to the manufacturer’s protocol. For primers used to amplify *Hbb-b1* and *Gapdh,* see **Table S2**. qPCR reactions were assembled using iQ SYBR Green Supermix (BioRad) and quantified on a Stratagene MX3000P qPCR machine. Expression fold changes were calculated using the ΔΔCt method.

#### Microscopy

Live cells were imaged in bright field on an Olympus CKX41 microscope.

#### RT-PCR after Pladienolide B treatment

For total RNA samples, RNA was extracted from approximately 5 million cells treated with Pladienolide B as described above and using TRIzol Reagent (ThermoFisher) according to the manufacturer’s protocol. For nascent RNA samples, RNA was extracted from the chromatin pellet after subcellular fraction as described above, except with the addition or not of Pladienolide B to all subcellular fractionation buffers at a final concentration of 1 μM. Poly(A)+ RNA was further depleted from this sample as described above. cDNA was generated from all RNA samples with SSIII RT (ThermoFisher) using random hexamer primers (ThermoFisher) according to the manufacturer’s protocol. PCR was performed using Phusion High-Fidelity DNA Polymerase (NEB) according to the manufacturer’s protocol. For the list of intron-flanking primers used in these experiments, see **Table S2**.

#### PacBio Sequencing Library Preparation

##### Genome-wide nascent RNA sequencing

Nascent RNA was isolated as described above from cells uninduced and treated with 2% DMSO for 5 days. A DNA adapter (**Table S2**) was ligated to 3’ ends of nascent RNA using the T4 RNA ligase kit (NEB) by mixing 50 pmol adapter with 300-600 ng nascent RNA. cDNA was generated from the adapter ligated RNA using the SMARTer PCR cDNA Synthesis Kit (Clontech), replacing the CDS Primer IIA with a custom primer complementary to the 3’ end adapter for first strand synthesis (**Table S2**). cDNA was amplified by 15 cycles of PCR using the Advantage 2 PCR Kit (Clontech), cleaned up using a 1x volume of AMPure beads (Agencourt), then PacBio library preparation was performed by the Yale Center for Genome Analysis using the SMRTbell Template Prep Kit 1.0 (Pacific Biosciences). The library was sequenced on four RSII flowcells and four Sequel 1 flowcells.

##### HBB targeted nascent RNA sequencing

Nascent RNA was isolated as described above from cells treated with 2% DMSO for 5 days, except that polyA(+) and ribosomal RNA depletion steps were omitted. A DNA adapter was ligated to 3’ ends as above, and custom RT primers were used to add barcodes during reverse transcription with SSIII reverse transcriptase (ThermoFisher; **Table S2**). cDNA was amplified by 26 cycles of PCR using the Advantage 2 PCR Kit (Clontech), but with custom gene-specific forward primers that were complementary to either a unique region in the 5’UTR of the human *HBB* gene or the endogenous mouse *Hbb-b1* gene in combination with the SMARTer IIA primer (Clontech). PCR amplicons were cleaned up with a 2X volume of AMPure beads (Agencourt), and PacBio library preparation was performed at the Icahn School of Medicine at Mt. Sinai Genomics Core Facility using the SMRTbell Template Prep Kit 1.0 (Pacific Biosciences). The library was sequenced on one Sequel 1 flowcell.

#### PRO-seq Library Preparation

##### Cell Permeabilization

All buffers were cooled on ice, all steps were performed on ice, and all samples were spun at 300 xg at 4°C unless otherwise noted. MEL cell differentiation was induced as previously described. Uninduced and induced cells were washed with PBS and resuspended in 1 ml Buffer W (10 mM Tris-HCl pH 8.0, 10 mM KCl, 250 mM sucrose, 5 mM MgCl_2_, 0.5 mM DTT, 10% glycerol), then strained through a 40 μm nylon mesh filter. 9X volume of Buffer P (Buffer W + 0.1% IGEPAL CA-630) was immediately added to each sample, cells were nutated for 2 minutes at room temperature, then spun for 4 minutes. Cells were washed in Buffer F (50 mM Tris-Cl pH 8.3, 40% glycerol, 5 mM MgCl_2_, 0.5 mM DTT, 1 μL/mL SUPERase.In [ThermoFisher]), then resuspended in Buffer F at a final volume of 1 x 10^6^ permeabilized cells per 40 μL. Samples were flash frozen in liquid nitrogen and stored at −80°C.

##### Library Generation

One million permeabilized uninduced and induced MEL cells were spiked with 5% permeabilized *Drosophila* S2 cells for data normalization and used as input for PRO-seq. Three biological replicates were generated per treatment condition. Nascent RNA was labeled through a biotin-NTP run-on: permeabilized cells was added to an equal volume of a 2X run-on reaction mix (10 mM Tris-HCl pH 8.0, 300 mM KCl, 1% Sarkosyl, 5 mM MgCl_2_, 1 mM DTT, 200 μM biotin-11-A/C/G/UTP (Perkin-Elmer), 0.8 U/μL SUPERase.In [ThermoFisher]), and incubated at 30°C for 5 mins. RNA was isolated using the Total RNA Purification Kit (Norgen Biotek Corp). Fragmentation of isolated RNA was performed by base hydrolysis with 0.25 N NaOH for 9 minutes on ice, followed by neutralization with 1X volume of 1 M Tris-HCl pH 6.8. To select for nascent RNAs, 48 μL of washed Streptavidin M-280 magnetic beads (ThermoFisher) in binding buffer (300 mM NaCl, 10 mM Tris-HCl pH 7.4, 0.1% Triton X-100) was added to the fragmented RNA, and samples were rotated at room temperature for 20 minutes. The Streptavidin M-280 magnetic beads were washed twice in each of the following three buffers: high salt buffer (2 M NaCl, 50 mM Tris-HCl pH 7.4, 0.5% Triton X-100), binding buffer (above), and low salt buffer (5 mM Tris-HCl pH 7.4, 0.1% Triton X-100). Beads were resuspended in TRIzol Reagent (ThermoFisher) and heated at 65°C for 5 mins twice to elute the RNA from the beads. A subsequent ethanol precipitation was performed for RNA purification. Nascent RNA was resuspended in 10 uM of the VRA3 3’ end adapter (**Table S2**). 3’ end ligation was performed using T4 RNA ligase I (NEB) for 2 hours at room temperature. A second Streptavidin M-280 magnetic bead binding was performed to enrich for ligated nascent RNAs. The beads were subsequently washed twice in high, binding, and low salt buffers, then once in 1X ThermoPol Buffer (NEB). To prepare nascent RNA for 5’ end adapter ligation, the 5’ ends of the RNA were decapped and repaired. 5’ end decapping was performed using RNA 5’ Pyrophosphohydrolase (NEB) at 37°C for 1 hour. The beads were washed once in high and low salt buffer, then once in 1X T4 PNK Reaction Buffer (NEB). Samples were treated with T4 Polynucleotide Kinase (NEB) for 1 hour at 37°C for 5’-hydroxyl repair. Next, T4 RNA ligase I (NEB) was used to ligate the reverse 5’ RNA adapter VRA5 (**Table S2**) as described previously. Following the 5’ end ligation, beads were washed twice in high, binding, and low salt buffers, then once in 0.25X FS Buffer (ThermoFisher). Reverse transcription was performed using Superscript IV Reverse Transcriptase (ThermoFisher) with 25 pmol of the Illumina TRU-seq RP1 Primer (**Table S2**). The RT product was eluted from the beads by heating the samples twice at 95°C for 30 seconds. All libraries were amplified by 12 cycles of PCR with 12.5 pmol of Illumina TRU-seq RPI-index primers, excess RP1 primer, and Phusion Polymerase (NEB). The amplified library was purified using the ProNex Size-Selective Purification System (Promega) and sequenced using NextSeq 500 machines in a mid-output 150 bp cycle run.

#### PRO-seq Data Preprocessing

Cutadapt was used to trim paired-end reads to 40 nt, removing adapter sequence and low quality 3’ ends, and discarding reads that were shorter than 20 nt (-m20 -q 1). Additionally, in order to align reads using Bowtie, 1 nt was removed from the 3’ end of all trimmed reads. Trimmed paired-end reads were first mapped to the *Drosophila* dm3 reference genome using Bowtie, and subsequent uniquely mapped reads to the dm3 genome were used to determine percent spike-in return across all samples. Paired-end reads that failed to align to the dm3 genome were mapped to the mm10 reference genome. Read alignment to the dm3 and mm10 genomes were performed with settings -k1 -v2 -best -X1000 --un. SAM files were sorted using samtools. Read pairs uniquely aligned to the mm10 genome were separated, and strand-specific single nucleotide bedGraphs of the 3’ end mapping positions, corresponding to the biotinylated RNA 3’ end, were generated. Due to the “forward/reverse” orientation of Illumina paired-end sequencing, “+” and stranded bedGraph files were switched at the end of the pipeline (Mahat et al., 2016). bedGraph files across replicates in each cell treatment were merged by summing the read counts per nucleotide position. Since the spike-in return was comparable between biological replicates within a treatment type, and no comparisons were made between the two treatment conditions, no further normalizations were performed.

#### PRO-seq and total RNA-seq Data Analysis

A list of active transcripts in MEL cells was first generated using PRO-seq signal within a 300 nt window around annotated TSSs in the GENCODE mm10 vM20 annotation. Intron annotations that did not correspond to an actively expressed transcript and had zero spliced read counts, suggesting no evidence of the intron’s usage in MEL cells, were removed. Additionally, if two intron annotations shared a 5’SS or 3’SS, the annotation with the most spliced reads was kept. Additionally, if introns shared both a 5’SS and 3’SS, the intron with the lowest annotated intron number was kept. Finally, first intron annotations were removed for Figure 4 metagene plot. For all other metagene analyses, introns within 750 nt of a TSS, and introns with fewer than 10 reads were also filtered out from the final list of unique introns to avoid bleed-through PRO-seq signal from the promoter-proximally paused Pol II. Metagene plots around the TSS, splice sites and PAS were generated by plotting the average PRO-seq reads (of three biological replicates) in uninduced cells at each indicated position with respect to the TSS, 5’SS, 3’SS or PAS respectively. Violin plots evaluating PRO-seq 3’ end or RNA-seq read coverage were generated by summing the signal at the indicated positions with respect to the 5’SS, 3’SS or PAS. P-values were calculated using either the Mann-Whitney or the Wilcoxon matched-pairs signed rank test

In order to extract PRO-seq reads that were spliced, filtered and trimmed PRO-seq reads were mapped to the mm10 reference index using STAR with the following changes to default settings: -- outMultimapperOrder Random --outFilterType BySJout-alignSJoverhangMin 8 --outFilterIntronMotifs RemoveNoncanonicalUnannotated. All reads in BAM format were filtered for reads that contained an “N” in their CIGAR string using pysam. Resulting reads were filtered to discard reads with an “N” size > 10,000 using pysam to remove poorly mapped reads or reads mapped across very large introns. In all analysis except where noted, PROs-seq data are represented as three biological replicates combined.

#### Long-read Sequencing Data Preprocessing

##### Genome-wide nascent RNA sequencing

Combined consensus sequence (CCS) reads were generated in FASTQ format, and Porechop was used to separate chimeric reads and trim external adapters with the SMRTer IIA sequences AAGCAGTGGTATCAACGCAGAGTAC and GTACTCTGCGTTGATACCACTGCTT with settings -- extra_end_trim 0 --extra_middle_trim_good_side 0 --extra_middle_trim_bad_side 0 -- min_split_read_size 100. Cutadapt was used to remove the unique 3’ end adapter on all reads in two rounds of filtering. First any reads with the adapter at the 3’ end were trimmed with settings -a CTGTAGGCACCATCAAT -e 0.1 -m 15 --untrimmed-output=untrimmed.fastq, and any reads which did not contain the full adapter were retained and their reverse complement was generated. Then, a second round of filtering with cutadapt using the settings -a CTGTAGGCACCATCAAT -e 0.1 -m 15 --discard-untrimmed was used to remove adapters from the reverse complement reads, and reads without the 3’ adapter were discarded. This ensures that each read contains a successfully ligated 3’ adapter which marks Pol II position, and since sequencing occurs in both forward and reverse orientations randomly, it places all reads in the correct 5’ to 3’ orientation. Reads from the two adapter trimming steps were combined into a single file, then Prinseq-lite was used to remove PCR duplicates with settings -derep 1. Prinseq-lite was used again to trim 6 non-templated nucleotides added at the 5’ end by the strand-switching reverse transcriptase and the 5 nt of the 3’ end adapter UMI with settings -trim_left 6 -trim_right 5. Reads were then mapped to the mm10 genome using minimap2 with settings -ax splice -uf -C5 -- secondary=no, and the resulting SAM files were converted to BAM and BED files for downstream analysis using samtools and bedtools. Reads overlapping the 7SK genomic region (chr9:78175302,78175633 in the mm10 genome) were filtered using samtools before all downstream analyses. Non-unique reads (reads with the same read name appearing more than once in SAM files) were removed. All data generated using Nanopore sequencing from Drexler *et al.* (GEO: GSE123191) were downloaded in FASTQ format and mapped to either the hg38 or dm6 genome using minimap2 with settings -ax splice -ut -k14, then converted to SAM, BAM, and BED formats as above, and non-unique reads were also removed. All data were visualized in and exported from IGV to generate genome browser figures. In all analysis except where noted, LRS data are represented as two biological and two technical replicates combined.

##### HBB targeted nascent RNA sequencing

Porechop was used on raw FASTQ reads to remove external adapters and separate chimeric reads with the common forward sequence and the SMRTer IIA reverse sequence GACGTGTGCTCTTCCGATCT and GTACTCTGCGTTGATACCACTGCTT (as well as the reverse complement sequences) with settings --extra_end_trim 0 --extra_middle_trim_good_side 0 --extra_middle_trim_bad_side 0 -- min_split_read_size 100 --middle_threshold 75. Reads were filtered and trimmed if they contained the 3’ end adapter as described above using the 3’ end adapter sequence plus the barcode sequence (**Table S2**). Prinseq was used to demultiplex and trim reads as above, then cleaned FASTQ files were mapped to a custom annotation of the integrated *HBB* locus, which is based on the GLOBE vector (Miccio et al., 2008).

#### PolyA Read Filtering

##### Genome-wide nascent RNA sequencing

Mapped reads in SAM format were filtered to remove reads that contained a polyA tail using a custom script (available on Github). Briefly, mapped reads that had soft-clipped bases at the 3’ end were discarded if the soft-clipped region of the read contained 4 or more A’s and the fraction of A’s was greater than 0.9. Similarly, reads with soft-clipped bases at the 5’ end (resulting from minus strand reads) containing at least 4 T’s and having a fraction of T’s greater than 0.9 were discarded.

##### HBB targeted nascent RNA sequencing

Additional parameters were added to the above criteria for removing polyA-containing reads from targeted data mapped to the *HBB* locus based on empirical observation. Since the *HBB* locus is integrated randomly in the MEL genome, long uncleaved transcripts that have coverage past the annotated *HBB* locus read into random genomic regions and cause long stretches of mismatched soft-clipped bases. A custom script was used to filter polyA-containing reads but retain uncleaved transcripts (available on Github). Briefly, reads were discarded if: they contained a fraction of A’s or T’s greater than 0.7 in the soft-clipped region that starts past the end of the *HBB* locus annotation; they contained a fraction of A’s or T’s greater than 0.7 and 4 or more A’s or T’s in the soft-clipped region starting within 50 nt of the annotated PAS; they contained a stretch of soft-clipped reads greater than 20 nt that starts within the annotated *HBB* gene. Uncleaved reads with long stretches of soft-clipped bases that passed this filtering were then recoded to contain a match in the CIGAR string downstream of the PAS in order to include these regions of the long-reads in coverage calculations.

##### Splicing Status Classification and Co-transcriptional Splicing Efficiency (CoSE) Calculation

The annotation of introns contained in active transcripts (described above for PRO-seq), was first filtered for unique intron start and end coordinates. Additionally, an upstream intron in *Hbb-b1* that was clearly not used in the LRS reads was removed (ENSMUST00000153218.1_intron_1_0_chr7_103827887_r). The resulting introns were extended by 1 nt on either end and were overlapped with bed files of long-reads using bedtools intersect in order to get regions of long-reads that spanned entire introns. Spliced junction coordinates in intron-overlapping long-reads were compared to the coordinates of each intron they overlapped to determine if the overlapped intron was spliced in the read. If the junction was not present in the read, a 10 nt window was included in the search for the junction to allow for slight mismatches in alignments. If the junction was not found, the intron was classified as unspliced. Next, reads which did not span the entire intron, but spanned at least 35 nt upstream of a 3’SS and were unspliced were counted toward the unspliced count for an intron. To classify splicing status of each read, the number of spliced introns was compared to the total number of introns that was overlapped. To calculate co-transcriptional splicing efficiency (CoSE), the splicing status classification of each intron was recorded as above, and the number of spliced introns and unspliced introns was summed per intron. Introns with identical 5’SS or 3’SS were filtered to keep only the intron with the most total reads. Introns with no spliced reads (no evidence of usage in MEL cells), introns that were longer than 10 kb, and introns covered by fewer than 10 reads were removed. For the remaining introns, CoSE was calculated by dividing the number of spliced reads by the total number of reads.

##### Distance from Splice Junction to 3’ End Calculation

Splicing intermediates (defined below), were filtered out from the long-read data in this analysis, since their 3’ ends do not represent the position of Pol II, but rather an upstream exon between step I and step II of splicing. For all remaining reads, data in were filtered for reads that contained at least 1 splice junction, and then the last “block size”, which represents the distance from the most distal splice junction to the 3’ end of the read, was calculated. Coordinates of the last spliced intron were also recorded, and each intron was matched to a transcript and categorized by gene biotype using mygene in python. Introns were matching to their corresponding transcript expression level using PRO-seq TSS counts as described above. To determine if certain genes exhibited a longer or shorter distance from 3’ end to Pol II, the distance was split into three equal size categories and transcript IDs from each category were entered into the online PANTHER classification system: no significant enrichment was obtained.

##### Splicing Intermediates Analysis

Long-reads were categorized as being splicing intermediates if the 3’ end of the read aligned exactly at the −1 position of an intron (last nucleotide of an exon). Introns considered in this analysis were the same set of introns considered for CoSE as described above. The number of intermediates aligned upstream of each intron was counted using bedtools intersect. The Normalized Intermediate Count (NIC) was calculated for each intron which was covered by at least 10 reads (as above) by dividing the number of splicing intermediate reads by the sum of splicing intermediate reads and spliced reads. The sequence of the 23 nt region surrounding intron 3’SS (−20:+3) and the 9 nt region surrounding the 5’SS (−3:+6) were extracted using bedtools getfasta, and these sequences were used to calculate 5’ and 3’ splice site scores using MaxEntScan (Yeo and Burge, 2004).

##### Long-read Coverage

Transcript coordinates associated with active TSSs (as described above) were obtained from UCSC. Transcripts were then grouped by the parent Gene ID, and the largest range of start and end coordinates from the grouped transcripts was kept. Library depth was then calculated using bedtools coverage across this file of collapsed active gene coordinates. Metagene plots of 5’ end, 3’ end, and entire read coverage across the same gene coordinates were generated using deepTools. Briefly, coverage was calculated and normalized by RPKM using the bamCoverage function, then coverage was scaled over all genes using the computeMatrix scale-regions function, and plots were generated using the plotProfile function. For coverage downstream of the PAS, long-reads were separated by splicing status (see below), then coverage was calucated using bedtools within a window around PASs that corresponded to active TSSs or specifically to a window around the *HBB* PAS. Coverage at all positions was normalized to the coverage at the position 100 nt upstream of the PAS. For coverage of splicing intermediates, bedtools coverage was used to calculate coverage of 5’ ends and 3’ ends across a 50-nt window around 5’SS and 3’SS of introns contained in active transcripts.

##### Uncleaved Transcripts Analysis

Bedtools intersect was used to identify long-reads with 5’ ends originating in a gene body of active transcripts (as described above). Reads were then categorized as being uncleaved transcripts if their 3’ ends were greater than 50 nt downstream of the PAS of the gene which the 5’ end overlapped with. In the case where a read 5’ end intersected multiple overlapping transcripts, it was only assigned as an uncleaved read if the 3’ end was downstream of all transcript PASs. Splicing status classification of uncleaved transcripts was carried out as described above.

##### HBB^IVS-110(G>A)^ Splicing and 3’ End Cleavage Analysis

For long-reads derived from HBB^IVS-110(G>A)^ cells, only reads that were spliced at intron 1 using the cryptic splice site were analyzed, and the rare reads with a splice junction using the canonical splice site were discarded. Splicing status classification, counting of splicing intermediates, and calculating coverage downstream of the PAS were performed as described above but with the custom *HBB* annotation coordinates.

## QUANTIFICATION AND STATISTICAL ANALYSIS

All information about statistical testing for individual experiments can be found in figure legends, including statistical tests used, number of replicates, and number of observations.

**Table S1.**
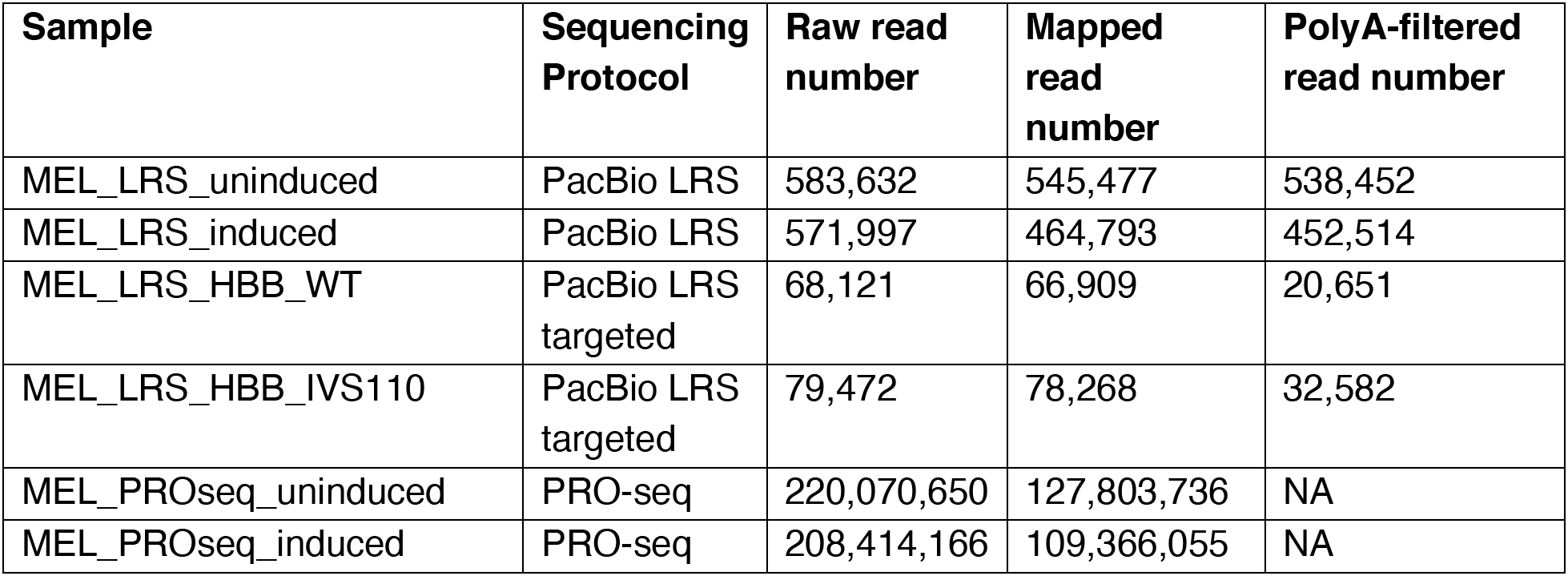
Related to Figure 1. RNA sequencing and mapping statistics. Read counts representing raw reads, mapped reads, and polyA-filtered reads (where applicable) for genome-wide LRS, *HBB*-targeted LRS, and PRO-seq. LRS samples represent combined counts for two biological and two technical replicates in each induction conditions, and PRO-seq samples represent combined counts for three biological replicates in each induction condition.

**Table S2.**
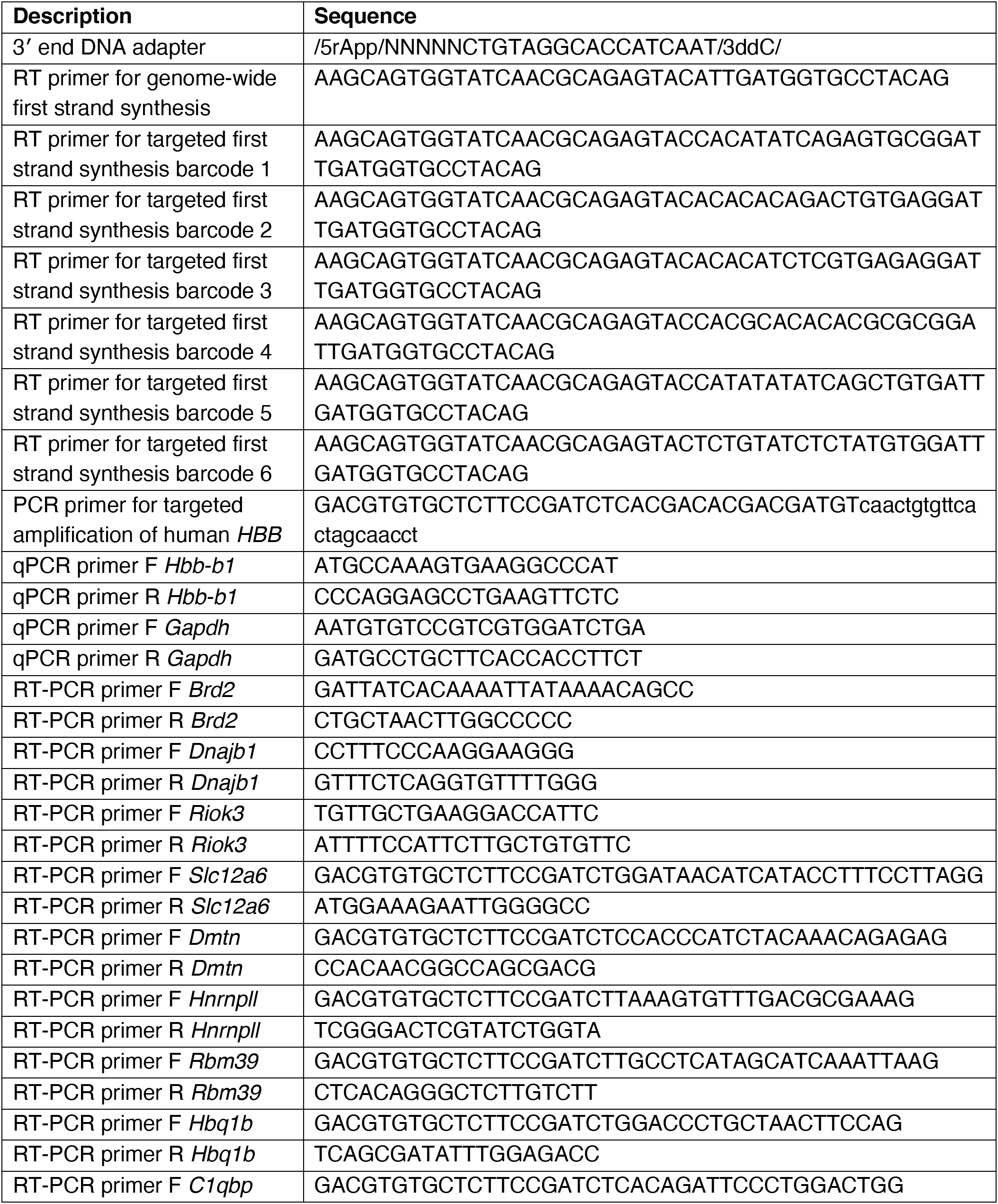
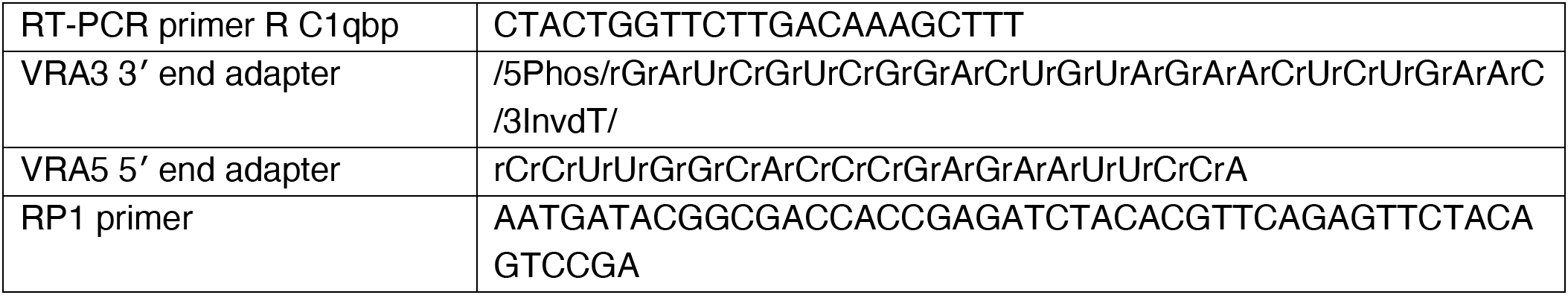
Related to STAR Methods. Oligonucleotides used in this study. Oligonucleotide sequences used in this study for LRS library preparation, qPCR, RT-PCR, and PRO-seq library preparation.

## KEY RESOURCES TABLE

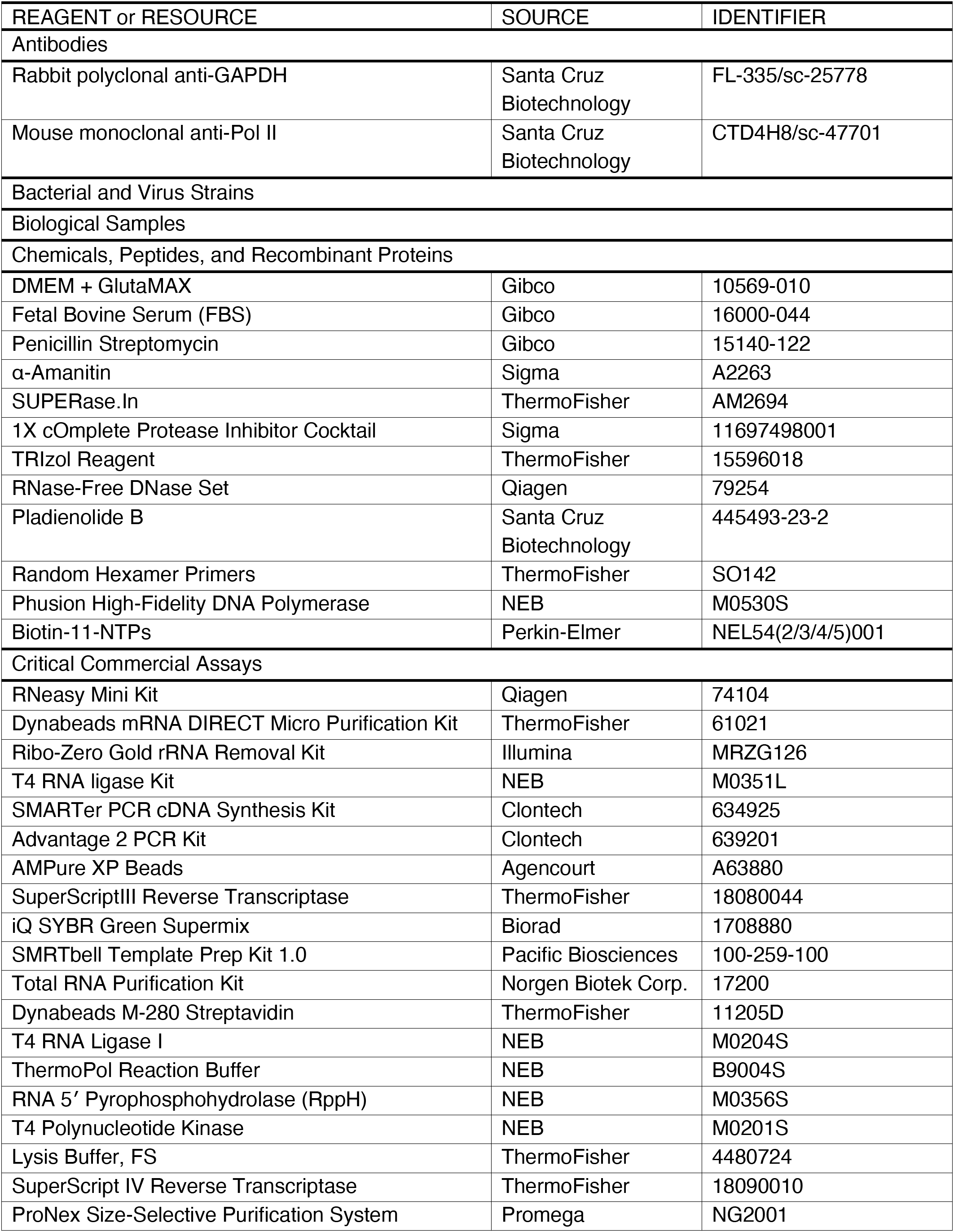
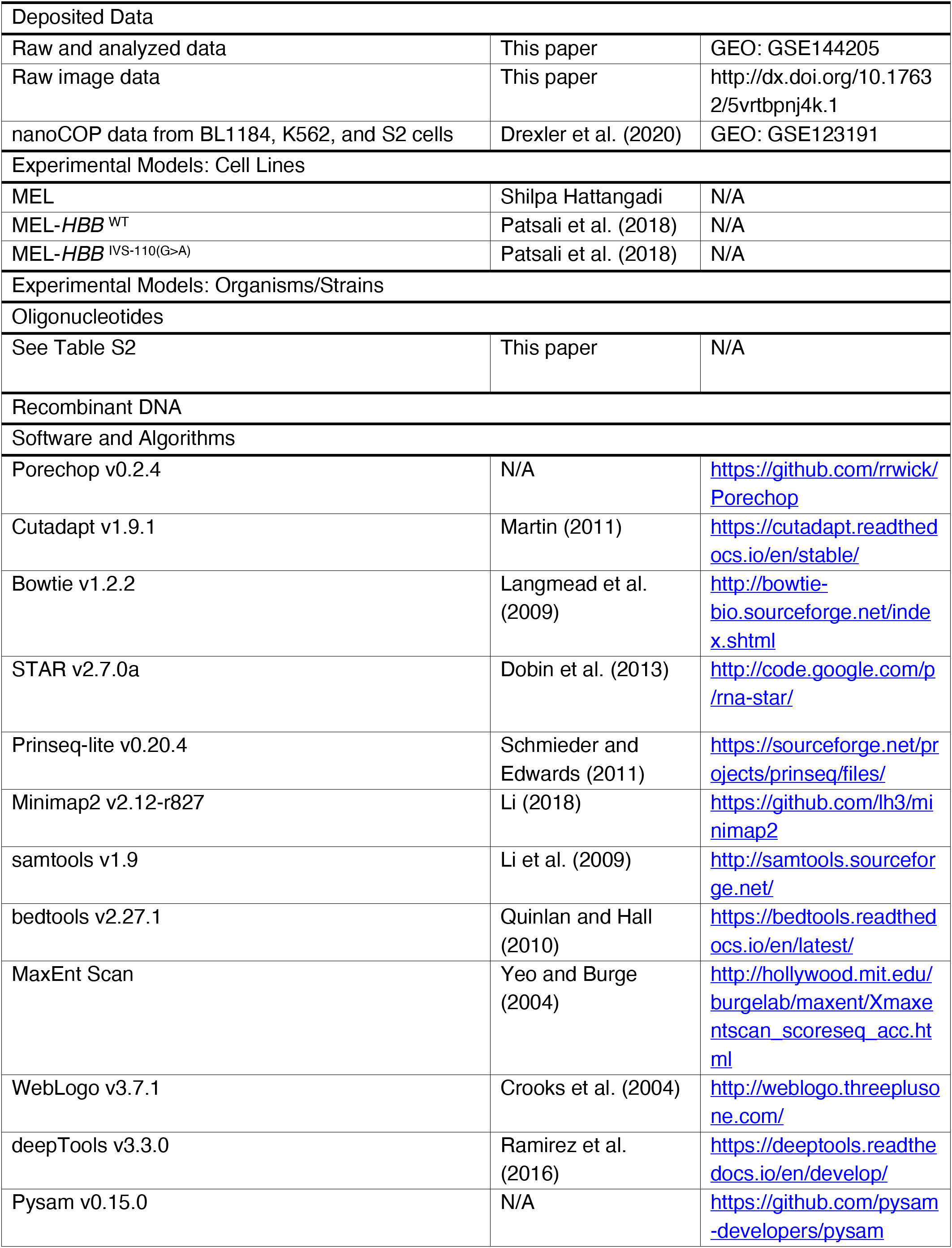
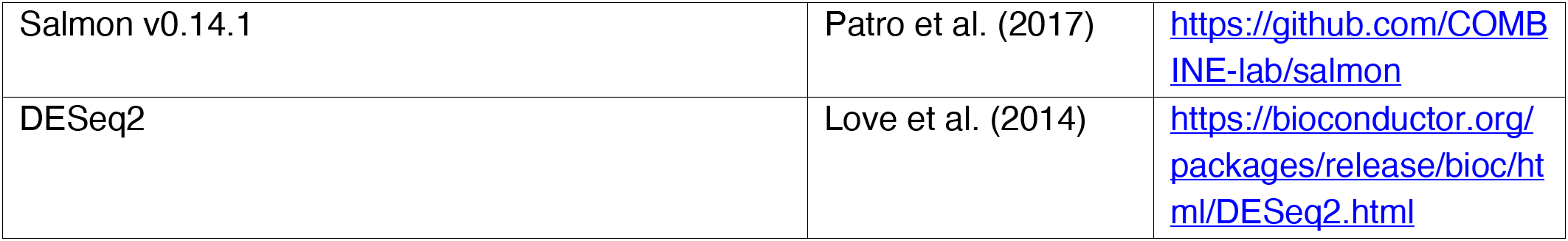

